# The curriculum effect in visual learning: the role of readout dimensionality

**DOI:** 10.1101/2025.09.19.677409

**Authors:** Charlotte Volk, Christopher C. Pack, Shahab Bakhtiari

## Abstract

Generalization of visual perceptual learning (VPL) to unseen conditions varies across tasks. Previous work suggests that training curriculum may be integral to generalization, yet a theoretical explanation is lacking. We propose an explanatory theory of visual learning generalization and curriculum effects by leveraging an artificial neural network (ANN) model of VPL in comparison with humans. We found that easy-to-hard sequential training improved generalization in both humans and ANNs. However, when easy and hard conditions were interleaved, humans and ANNs showed different behaviours: while ANNs performed worse than with sequential training, humans maintained good performance but with large inter-individual variability. Investigating ANN models trained with different curricula, we demonstrated that models relying on low-dimensional neural populations showed superior generalization. This *readout subspace dimensionality* was directly determined by curriculum: learners who learned from easy tasks early formed lower-dimensional subspaces and generalized better. Our theory provides a mechanistic framework linking curriculum design to VPL generalization through neural population dimensionality.

**Author Summary:** Learning new skills is fundamental to humans and animals. However, a key challenge in learning is generalization: applying learned skills to new situations not encountered during training. While it is well known that training curriculum affects how well we generalize, the underlying mechanisms remain poorly understood. What makes certain training curricula more effective than others? Why do some learners generalize better than others even with similar training? We developed a computational theory to explain how curriculum design influences generalization in visual learning. Using a combination of human behavioural experiments and artificial neural network modeling, we posit that easy-to-hard training sequences lead learners to focus on fewer, more essential visual features. This creates low-dimensional neural representations that are robust and generalize well to new conditions. Importantly, we found this benefit even extends to learners who spontaneously adopt easy-to-hard strategies on their own, without explicit curriculum design. Our theory provides a mechanistic link between curriculum structure, neural population dynamics, and generalization performance. Moreover, these findings offer practical guidelines for designing effective training programs in education and rehabilitation.

## Introduction

Visual perceptual learning (VPL) enables humans to enhance their visual perception through practice [1]. However, a fundamental limitation of VPL is that the improvements often fail to generalize beyond the trained conditions [2,3]. The ability to transfer improvements to new conditions is known as generalization [4], and varies considerably across visual tasks [3,5]. While some training protocols and tasks lead to better generalization, other training produces highly specialized improvements that do not transfer even to slightly different conditions [3,5]. This limited generalization substantially constrains the practical applications of VPL in real-world settings, motivating extensive research into understanding and overcoming these constraints.

A particularly important observation has been the sequential effect of training tasks on learning generalization. The seminal work of Ahissar and Hochstein [3] demonstrated that more difficult visual tasks lead to lower generalization. In particular, they showed that brief exposure to an easy orientation discrimination task dramatically improved subsequent generalization in a more difficult version of the same task. Recent studies have further shown that initial training on more generalizable tasks (i.e., tasks that often lead to better learning transfer) can improve generalization on subsequent tasks [6,7]. Beyond strictly sequential paradigms, studies have shown that interleaving different task conditions can also enhance generalization [8,9]. These findings suggest that the order and structure of training can substantially impact learning outcomes, but it is unclear in which scenarios a sequential curriculum would be advantageous over an interleaved curriculum.

Despite substantial experimental evidence supporting the efficacy of certain training curricula in enhancing generalization, our theoretical understanding of why and how specific training curricula may influence learning generalization remains limited. Existing theories explain potential causes of variability in generalization across tasks [10] but provide little insight into how the sequential order of tasks affects generalization. A theoretical formulation of this phenomenon is important for a better understanding of the neural and computational basis of learning in the brain, but also crucial for developing optimal learning curricula that maximize generalization in any real-world application of VPL.

In this paper, we hypothesize that the effectiveness of different learning curricula depends on how many distinct visual features the brain recruits during training to solve the task. Specifically, we predict that curricula beginning with easier tasks will lead learners to rely on fewer, more essential visual features, resulting in better generalization to unseen conditions. For example, in learning to discriminate between rotated images, easy training (large rotation differences) might lead the visual system to discover that edge orientation alone is sufficient for discrimination. However, hard training (small rotation differences) might recruit more additional features (e.g., texture patterns, local contrast, multiple orientation channels, etc.) to achieve higher accuracy. While this high-dimensional multi-feature solution works for trained conditions, it becomes brittle when some of the features are absent in new contexts (i.e., test condition). In contrast, the simple low-dimensional edge-orientation solution remains robust across diverse conditions. This framework predicts that easy-to-hard curricula will produce more generalizable learning by constraining the visual system to discover low-dimensional, feature-sparse solutions that capture the essential structure of the task.

We employed artificial neural networks (ANNs) and psychophysics to test this hypothesis and develop theoretical insights into the sequential and interleaved learning paradigms. ANNs have proven to be valuable models of biological visual systems [11–14], capable of predicting various neural and behavioural aspects of visual processing [15,16]. Moreover, Wenliang and Seitz [17] demonstrated that an ANN model of the ventral visual pathway can reproduce key VPL phenomena when fine-tuned for orientation discrimination tasks. We used a modified version of their model to investigate the effects of learning curricula on generalization. Specifically, we compared three training approaches: (1) a sequential curriculum where an easy, generalizable task condition preceded a hard, non-generalizable task condition; (2) a shuffled curriculum where easy and hard task conditions were randomly interleaved; and (3) a baseline non-sequential paradigm focused solely on the hard task condition.

We conducted parallel experiments with both human observers and ANNs using similar training curricula on an orientation discrimination task. By comparing learning dynamics between the observers and ANNs, we identified the similarities and the dissimilarities between human observers and the artificial learning systems. Our comparisons confirmed that the sequential curriculum improved generalization compared to non-sequential training in both humans and ANNs. Interestingly, however, humans exhibited similar average generalization improvements across both sequential and shuffled conditions, while ANNs benefited more from explicit sequential ordering of easy to hard tasks. Closer inspection revealed a sequential easy-to-hard “implicit curriculum” in humans and ANNs in the shuffled condition, although there was substantial variability in the strength of this curriculum across humans. We found that the strength of the implicit curriculum correlated with the generalization performance in observers, and that forcing ANNs to mimic the learning dynamics of humans in the shuffled condition reproduced this correlation.

By examining the model’s readout subspace – the specific neurons and visual features that contribute to the network’s decisions – we found strong support for our hypothesis. Learning generalization had a strong negative correlation with how many neurons with distinct visual feature preferences contributed more strongly to the readout, with networks using fewer visual features, or lower-dimensional readouts, showing better transfer to new conditions. Critically, this feature recruitment pattern was largely determined during early training steps, when the network first learned which neural dimensions and features were most informative. This finding provides a mechanistic explanation for why easy-to-hard curricula improve generalization: initial training on easier tasks constrains the network to discover simpler, more essential feature combinations that remain robust across diverse testing conditions (*3, 7*). Our results thus establish a direct link between curriculum design and generalization capacity through the dimensionality of the readout subspace.

## Results

### Visual learning task and the ANN models

We studied visual learning in an orientation discrimination task in both convolutional ANN models (Fig 1A) and human observers (Fig 1B), where the models and observers were trained to classify the rotation direction of a Gabor stimulus relative to a reference orientation. During training, the reference orientation was 145°, and the stimuli could occur in two diagonally opposite positions on the screen, in the upper-left and bottom-right corners. To evaluate generalization in the human observers, we swapped the diagonal that stimuli were presented on to the upper-right and bottom-left corners of the screen and changed the reference orientation of the stimuli during the test to 55° (Fig 1C) [18]. However, testing on a different location in the visual field may not cause specificity in convolutional ANNs, as they may be equivariant to spatial location. To evaluate generalization in the models, we used a Gabor stimulus which maintained the training angle separation but with doubled spatial frequency. Separately, we also evaluated generalization for a different reference angle orientation to confirm the robustness of our results (S2 Fig). We did not use spatial frequency as a transfer condition for the human observers to avoid potential variability induced due to spatial frequency preferences across observers [19,20].

**Fig 1.**
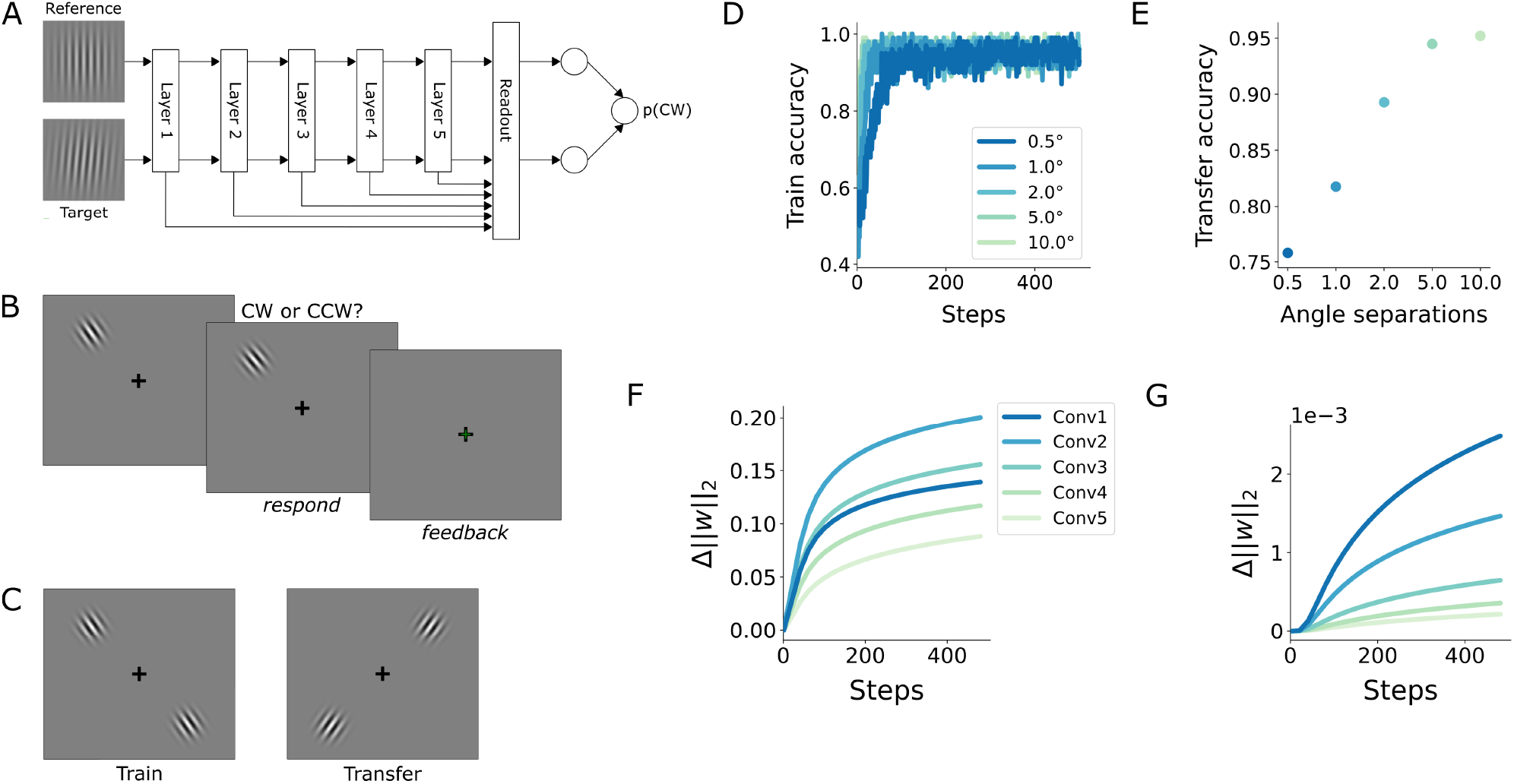
An ANN model of visual learning. (**A**) Schematic of the ANN model of the ventral visual pathway. The network was a convolutional ANN (based on AlexNet) pretrained on object categorization. (**B**) Schematic of the experimental task. Observers viewed a reference image, target image, and then provided a keypress response. Observers were given feedback during training. (**C**) Left: Train condition. Stimuli were presented in the upper-left or bottom-right corner, randomly chosen in each trial. The reference orientation was 145°. Right: Transfer condition. Stimuli were presented in the upper-right or bottom-left corner, with a 55° reference orientation. (**D**) Train accuracy throughout training for models trained on a single angle separation. (**E**) Transfer accuracy after training. This pattern of generalization (i.e., worse generalization for smaller angle separations) followed the pattern previously observed in humans. (**F**) Change in the L2-norm of skip connection weights (connecting convolutional layers to the output) throughout training on a 0.5° angle separation. (**G**) Change in the L2-norm of convolutional weights throughout training on a 0.5° angle separation. Note the difference in scale between (**F**) and (**G**). Error bars are not visible.

We used a convolutional ANN model based on AlexNet [21] and pretrained on ImageNet [22] that has demonstrated strong alignment with both monkey [23] and human [24] visual systems and has been used to replicate several behavioural and neuronal attributes of visual learning [17]. Despite its relatively modest size and straightforward architecture, this model performs comparably to more sophisticated, larger architectures in terms of model-brain alignment [24]. Moreover, its relatively small size enables more thorough investigation of both its learning dynamics and post-training representations. We modified the model by incorporating readout weights, or “skip connections”, from every layer of the model to the output decision neuron (Fig 1A). These skip connections enable all hierarchical levels of the model to directly influence the final output, unlike pure feedforward networks where only the final processing stage directly contributes to the output [25]. This is consistent with neurophysiological evidence suggesting that visual areas across different processing stages can directly influence behavioural responses [26,27]. The modified model was fine-tuned on the orientation discrimination task using backpropagation.

First, our ANN model successfully reproduced the main behavioural characteristics of visual learning [3]. Accuracies increased more rapidly during training for tasks with larger angle separations (easier tasks) compared to those with smaller angle separations (harder tasks) (Figs 1D, S1A, S2A, S3A). Moreover, our model replicated the generalization pattern previously observed in humans [3], namely, lower generalization to a transfer condition for smaller angle separations (Figs 1E, S2B, S3B). These results are in general agreement with the previous findings of Wenliang and Seitz [17].

To identify the loci of plasticity underlying learning, we further examined changes in the model’s weights throughout the training process. Our analysis showed that the changes in the L2-norm of the weights in the skip connections (Figs 1F and S3C) or in the gradients of the weight norms (S1D Fig) were approximately 200x larger than those in the between-layer convolutional weights (Figs 1G, S3D, S1E). This observation suggests that learning predominantly occurred in the readout weights, consistent with previous neurophysiological studies indicating that VPL mostly alters the readout of sensory areas rather than their tuning properties [26,28,29]. This observation supports the hypothesis that direct skip connections to downstream association areas channel learning primarily into the readout weights from visual areas, rather than into the sensory representations of stimuli themselves [30]. To confirm this mechanism, we tested a variant of the model where all between-layer connection weights were frozen, allowing changes only in the readout connections. This restricted model exhibited identical behavioural characteristics and generalization patterns (S1B-C Fig). Therefore, our subsequent analyses of the model’s connection weights throughout this paper focus exclusively on the learned readout weights.

### The effect of training curricula on ANN models and humans

We next investigated how different training curricula affect generalization in a task that typically exhibits poor transfer (i.e., orientation discrimination with small angle separation). We trained both our ANN models and human observers using three distinct approaches: 1) a baseline non-sequential paradigm focused only on the harder version of the task; 2) a shuffled curriculum with randomized difficulty levels; and 3) a sequential easy-to-hard curriculum (Fig 2A). In the sequential curriculum, the models or observers were trained first on an easier version of the task (5° angle separation) for the first half of training, and then a harder version (1° angle separation) for the second half of training. In the shuffled curriculum, we trained on randomly interleaved 5° and 1° angle separations for the entire training process. The baseline non-sequential condition involved training exclusively on the 1° angle separation for an equivalent duration.

**Fig 2.**
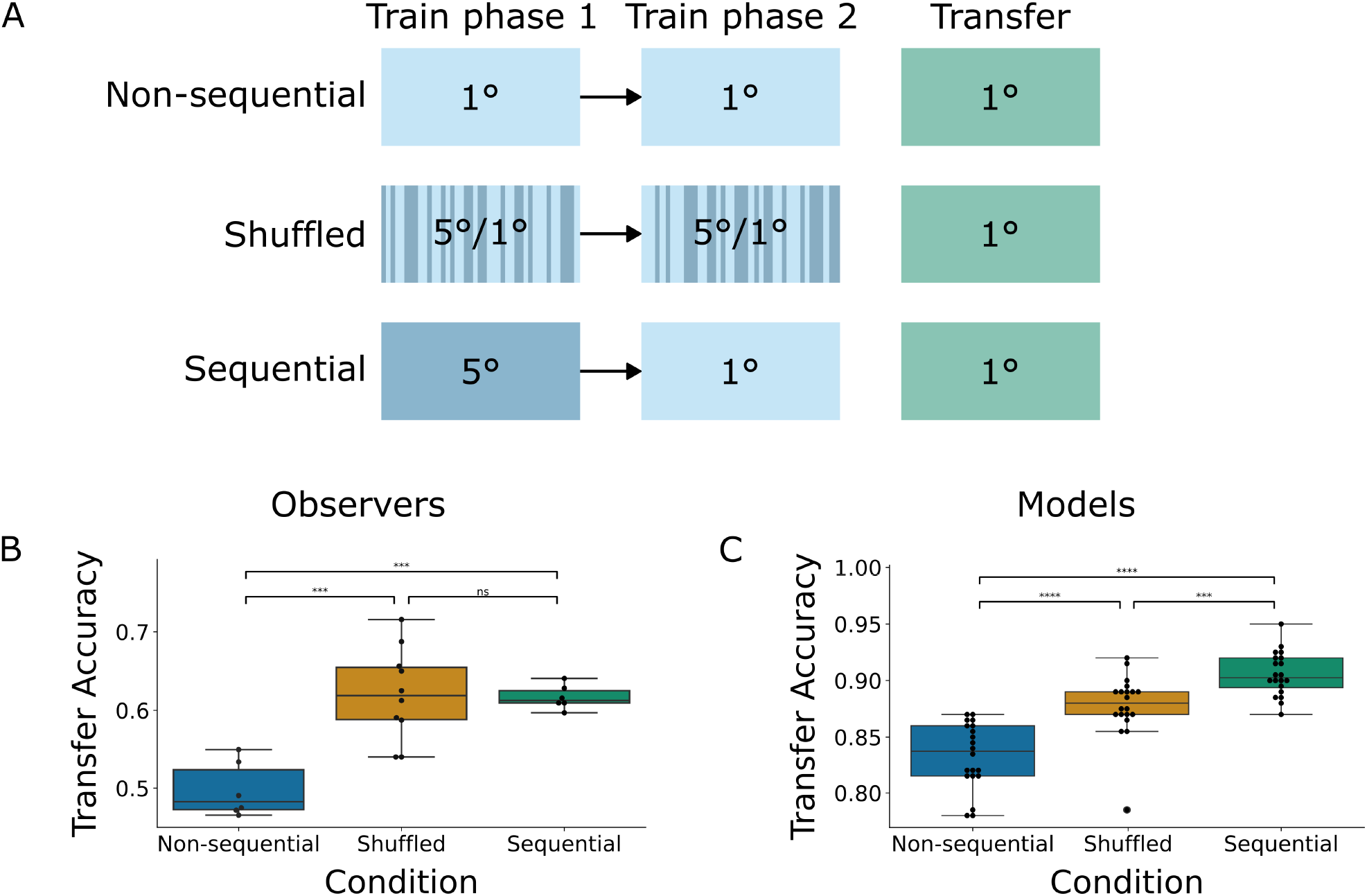
The effects of training curricula across models and observers. (**A**) Visualization of curricula that were used to train models and human observers. Dark blue indicates training on 5° angle separations, light blue indicates training on 1° angle separations, and green indicates a transfer condition on 1° angle separations. In a non-sequential curriculum, models and observers are trained only on 1° angle separations; in the shuffled curriculum, they are trained on randomly interleaved 5° and 1° angle separations; and in the sequential curriculum, they are trained for the first half of training on 5° and for the second half of training on 1° angle separations. (**B**) Transfer accuracy for human observers across curricula. Observers trained on either the shuffled or sequential curricula significantly outperformed observers trained on the non-sequential curriculum, and there was no significant difference between the shuffled and sequential curriculum. (**C**) Transfer accuracy for models across curricula. Models trained on either the shuffled or sequential curricula significantly outperformed those trained on the non-sequential curriculum, but models trained on the sequential curriculum also significantly outperformed those trained on the shuffled curriculum.

Our results showed that the shuffled and sequential curricula significantly improved generalization compared to the baseline non-sequential condition in both human observers (Fig 2B; shuffled: *p*=0.00013; sequential: *p*=0.00016) and the ANN model (Figs 2C, S2C, S3E; shuffled: *p*<<0.0001; sequential: *p*<<0.0001). However, while the sequential curriculum significantly outperformed the shuffled curriculum in the model (*p*=0.0008), interestingly, human observers showed comparable improvement with both curricula, with no significant performance difference between them (*p*=0.84). Notably, the shuffled condition in human observers displayed much larger variance than the sequential condition, suggesting substantial individual variability. Further analysis of the learning dynamics provided insight into the discrepancy between the models and observers in the shuffled condition.

### Implicit curricula in the shuffled condition predict generalization in ANN models and human observers

We noticed that the human observers in the shuffled condition had substantial variability in their performance relative to those in the sequential condition. We wanted to investigate the source of this variability and additionally attempt to determine the reason for the discrepancy between the performance on the shuffled condition between the human observers and the model.

We first found that observers trained on the shuffled condition appeared to follow an “implicit curriculum”: observers plateaued at approximately 90% accuracy on the easy condition about halfway through training but continued to learn from the hard condition until the end of training (Fig 3A). This suggests that when given randomly interleaved tasks, human observers develop their own implicit easy-to-hard curriculum, consistent with previous observations by Ahissar and Hochstein [3].

**Fig 3.**
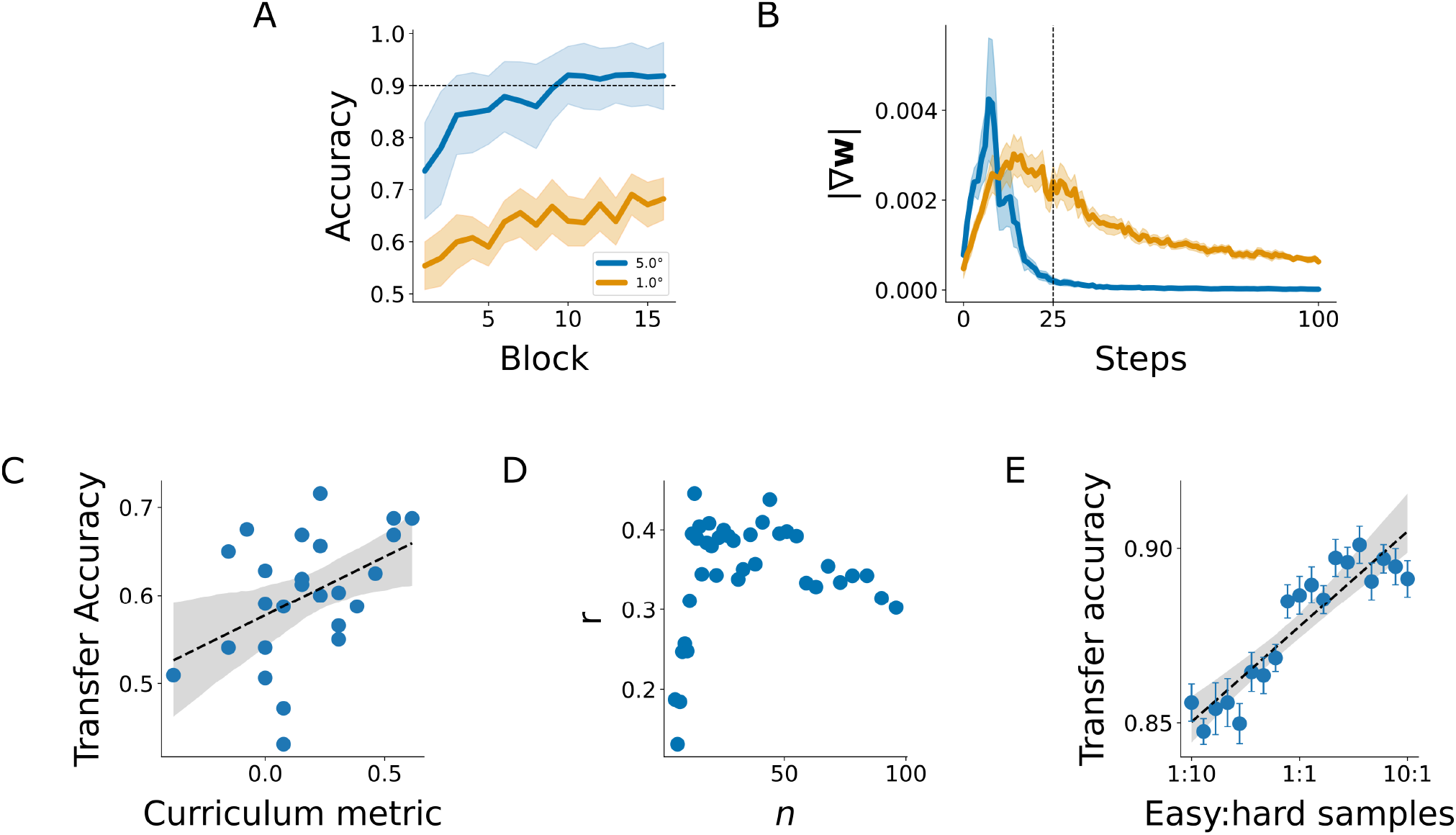
An implicit curriculum arises in human observers and models through shuffled training. (**A**) Accuracy throughout training for human observers on the shuffled condition shown separately for easy (5°) and hard (1°) samples. Accuracy on easy samples plateaued at around 90% starting approximately halfway through training, while accuracy on hard samples continued to improve. (**B**) Parameter gradients for models trained on the shuffled task condition in the first 100 steps of training. Parameter changes occurred predominantly in the easy condition (5°) early in training, then transitioned to greater changes in the hard condition (1°). (**C**) Transfer accuracy correlated with our curriculum metric across human observers – peak value from panel (D) (*r*=0.45, *p*=0.026). Our curriculum metric provided a proxy measure for the strength of the easy-to-hard implicit curriculum that the observer followed in the first *n* easy trials and the first *n* hard trials. (**D**) The correlation of transfer accuracy vs. curriculum metric, with varied *n* (number of trials used to calculate the curriculum metric). *n* was sampled logarithmically from the range [5, 100]. The correlation was robust for various values of *n* but decreased for very small and very large *n*. (**E**) Transfer accuracy of the models trained on a shuffled condition when different ratios of easy:hard samples were shuffled in for the first 25 steps of training (*r*=0.91, *p*<<0.0001). Learning from a higher ratio of easy samples at the beginning of training correlated with a higher transfer accuracy. Shading on all plots represents 95% CI.

Having observed implicit curricula in human observers, we asked whether similar implicit curricula also emerged in the ANN model during shuffled training. Unlike with humans, we can directly measure how much each task condition (hard vs. easy samples) changes the model’s parameters throughout training. We computed the gradients of the model’s training loss with respect to the model weights, which provided the direct learning signal. We used the mean of the absolute gradient values as a measure of the learning signal that drives the changes in a model’s performance. In the shuffled curriculum, parameter gradients peaked earlier for easy samples, and later for hard samples (Figs 3B and S4A). We observed a similar pattern in the accuracy of each condition (S4B Fig). This revealed an implicit quasi-sequential curriculum in the model: the model learned from easy samples first and then shifted focus to harder samples as training progressed. This finding is consistent with previous work showing that ANNs implicitly learn tasks in a highly consistent order [31,32]. However, the precise dynamics of this implicit curriculum may differ among human observers and between humans and ANNs. While ANNs learn the task in a very consistent order with little variability among model instances, human observers show greater variability in their learning dynamics. We therefore investigated this variability and its relationship with learning generalization.

We defined a “curriculum metric” (see Eq. 1 in Methods) as the mean accuracy on the first *n* easy trials minus the mean accuracy on the first *n* hard trials, providing a proxy measure of each observer’s implicit curriculum. We found a positive correlation (*r*=0.45, *p*=0.026) between the transfer accuracy and our curriculum metric across observers (Fig 3C), indicating that the implicit curriculum is related to generalization: An observer who followed an easy-to-hard implicit curriculum was more likely to generalize better. Interestingly, we found an inverse-U relationship between the correlation value and the number of trials *n* used for calculating our curriculum metric (Fig 3D). For very small and very large numbers of trials, the measured implicit curriculum does not predict generalization. Performance in initial trials is affected by multiple factors that may not properly reflect observers’ implicit curriculum. Additionally, variability in performance in later training trials is also not predictive of learning generalization. To confirm that the relationship between the transfer accuracy and our curriculum metric is not merely a consequence of the random assignment of easy and hard trials early in training, we measured the correlation between the transfer accuracy and the ratio of easy to hard samples during the first *n* overall trials. We found no significant correlations (all *p*-values >0.05 for *n* sampled logarithmically from the range [5, 100]) indicating that the variability in performance is due to the observers own implicit curriculum, not the explicit random order of samples.

To more directly investigate whether the dynamics of the curriculum explain performance differences, we manipulated the ratio of sample difficulty at the beginning of model training to indirectly control the models’ implicit curriculum dynamics. Unlike humans, models learn tasks in a predictable and consistent order in the shuffled condition without much variation between model instances [31,32]. Therefore, we must manually introduce variability by explicitly manipulating the sample difficulty in order to force a human-like curriculum. Specifically, we repeated our shuffled model training as before, but instead of randomly selecting task difficulty conditions (i.e., angle separations) for each training step, we modified the ratio of easy:hard task difficulties for the first 25 steps of training. The number of steps was chosen because the models’ parameter gradients on the easy condition drop to approximately zero in 25 training steps (Fig 3B). This concept is similar to the idea of “pacing” in curriculum learning for ANNs, which controls the amount of training examples sampled from each difficulty as training progresses [33,34].

We found a very strong positive correlation (*r*=0.91, *p*<<0.0001) between the ratio of easy:hard samples and the transfer accuracy (Fig 3E), replicating our findings in the human observers (Fig 3C). Additionally, models trained on ratios with a higher proportion of easy to hard samples performed better than a pure shuffled curriculum (1:1 ratio) (Fig 3E). This suggests that forcing a “more sequential” curriculum in the shuffled condition improves generalization, and the implicit curriculum that the model discovers through gradient descent-based learning (Fig 3B) is perhaps sub-optimal for the task.

### Readout subspace dimensionality explains generalization variability across curricula

Most learning in our model occurred in the readout weights that connected different network layers to the output (Figs 1F-G, S1D-E and S3C-D). Throughout training, each neuron received readout weights that ultimately determined its contribution to the model’s output. The neurons with the largest contributions and their visual feature preferences form the representational space that determines the model’s behaviour. We hypothesized that generalization depends primarily on the dimensionality of this feature space – specifically, that learning curricula establishing low-dimensional readout subspaces (recruiting fewer distinct visual features) result in better generalization. We tested this hypothesis by analyzing the readout weights of our ANN model after training.

First, after applying dimensionality reduction (Principal Component Analysis) on the readout weight vectors, we observed that readout weights followed distinct optimization trajectories for each curriculum throughout learning (Fig 4A), indicating differences in the final readout weight patterns across the training paradigms. These readout weights determine neuronal contributions across different layers to the output (the readout subspace). Although all models, regardless of their training paradigm (non-sequential, sequential, or shuffled), were performing the same task of small-angle discrimination at the end of their training, they read out from distinct subspaces of the representational space to solve the task due to their different training histories. These distinct readout subspaces underlie the variability in generalization across learning curricula.

**Fig 4.**
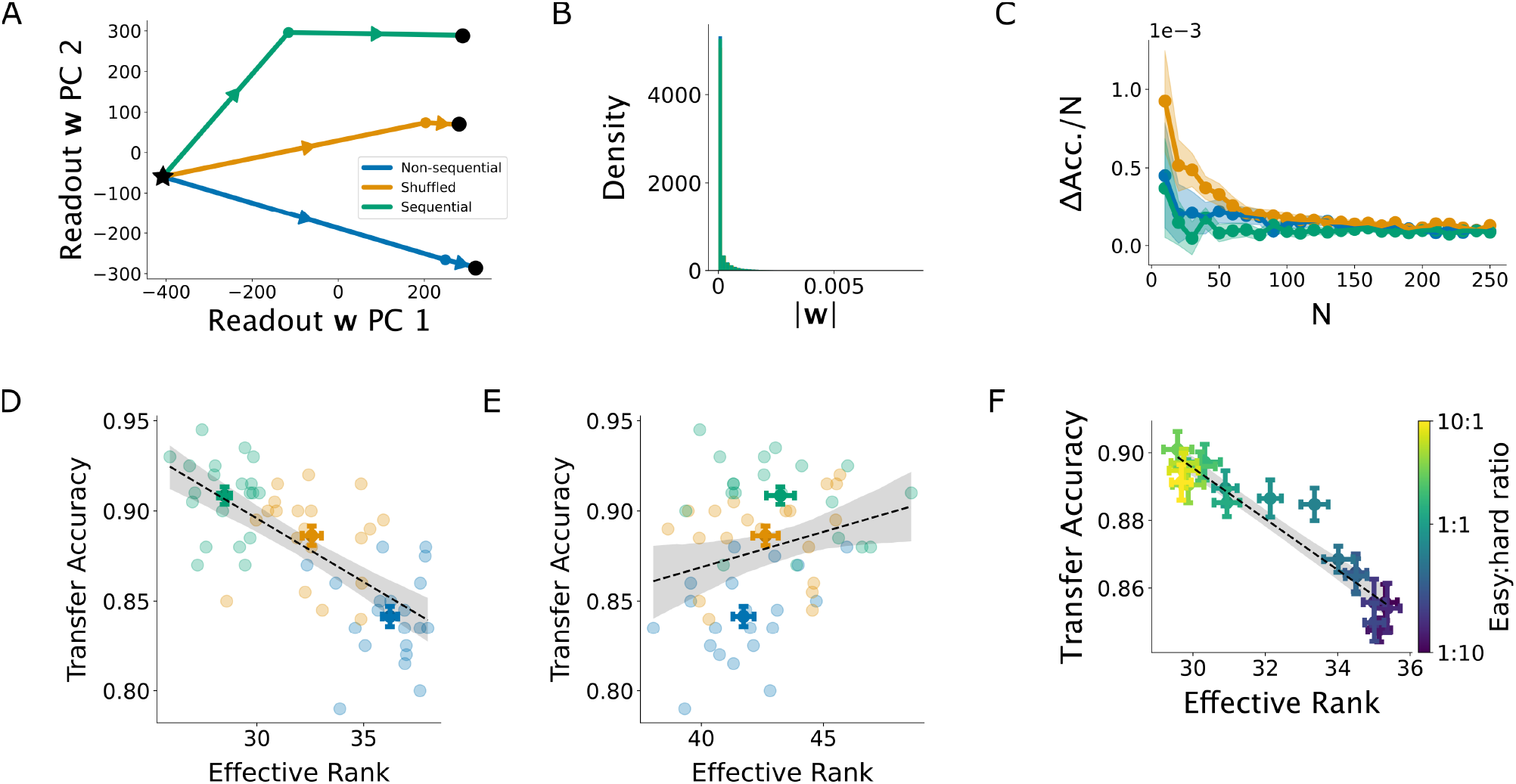
Readout subspace dimensionality determines generalization. (**A**) Projection of model readout weights onto their first two principal components for all three curricula at the beginning (marked by a black star), middle, and end (marked by black circles) of training. Arrows show the direction of training progression in weight space; different curricula follow different trajectories. (**B**) Histogram of all non-zero readout weights at the end of training. The majority of weights were zero, with only a small portion of non-zero weights. Distributions for the three curricula visually appear to overlap almost completely, and standard error is not visible. (**C**) Accuracy drop per neuron lesioned, plotted against the number of neurons lesioned. The accuracy drop per neuron plateaued at about N=150, meaning that the readout subspace was mostly affected by about 150 neurons. (**D**) Dimensionality of the 150-neuron readout subspace, measured by the effective rank, for non-sequential, sequential, and shuffled curricula (*r*=-0.69, *p*<<0.0001). Curricula leading to a low-dimensional readout subspace had a higher transfer accuracy. Results for other subspace sizes are shown in S5 Fig. **(E)** Dimensionality of the subspace created by sampling 150 random neurons from the readout. There is no apparent correlation in transfer accuracy and effective rank (*r*=0.25, *p*=0.058). (**F**) Negative correlation of the dimensionality of the readout subspace for shuffled curricula with different easy:hard sample ratios with transfer accuracy (*r*=-0.95, *p*<<0.0001). Models which were forced to learn from easy samples early in training had lower-dimensional readout subspaces and exhibited better generalization.

To further characterize these readout subspaces, we examined the distribution patterns of the learned weights. Analysis of learned readout weight distributions revealed high skewness in all models, with only a small portion of non-zero weights (Fig 4B). This suggests that a small subset of neurons, with larger readout weights, mainly determined the readout subspace of each model. To identify these “influential” neurons, we lesioned the neurons with the largest absolute readout weights in increments by removing their weights to the readout and quantified the effect of this lesioning on transfer accuracy. We found that the contribution to transfer accuracy per neuron asymptotically reached zero after approximately 150 neurons were lesioned, indicating that ∼150 influential neurons determined the model’s readout subspace (Fig 4C). However, our results are invariant to a range of subspace sizes (S5 Fig). We also repeated this analysis by choosing 150 neurons with the largest individual functional contribution to the transfer accuracy to confirm that our results are unaffected (S6 Fig).

Given these influential neurons, we could test our hypothesis: how many independent visual feature dimensions exist in the readout subspace of each model? In our non-stochastic model, this dimensionality reflects the number of neurons with dissimilar visual selectivity (different orientations, spatial frequencies, textures, etc.) that collectively form the model’s readout subspace. Larger numbers of neurons with diverse tuning properties contributing to the readout create higher-dimensional readout subspaces. According to our hypothesis, the observed generalization variability across training curricula stems from differences in readout subspace dimensionality.

To test this hypothesis, we estimated the dimensionality of readout subspaces across our three base curricula. From each model, we collected the activations of the 150 influential neurons with the largest absolute weight values in response to 100 natural images [22]. We estimated the dimensionality of each readout subspace using the effective rank [35,36]. We found that the effective ranks were negatively correlated with the transfer accuracies of the models – i.e., more generalizable readout subspaces also occupy a lower-dimensional representational space (*r*=-0.69, *p*<<0.0001) (Figs 4D, S2D, S3F, S6B). When we repeated this analysis for 150 randomly chosen neurons, we did not see a correlation between transfer accuracy and effective rank between curricula (*r*=0.25, *p*=0.058) (Fig 4E). This confirms that the dimensionality of the readout subspace is what differs here, not the overall dimensionality of the readout.

We also measured the effective rank of models with different imposed easy:hard sample ratios for the first 25 steps of training, finding a strong negative correlation between effective rank and transfer accuracy (*r*=- 0.95, *p*<<0.0001) (Figs 4F, S2E, S3G, S6C). A higher easy:hard ratio also led to a lower effective rank, corroborating the relationship between curriculum and dimensionality. This reinforces our main theoretical claim: the curriculum that leads to the lowest readout subspace dimensionality (i.e., relies on a smaller number of independent feature dimensions) leads to the highest generalization. In scenarios with limited observations (as in most VPL paradigms), relying on fewer essential visual features is more robust in transfer conditions than relying on many features, some of which may not be useful under unseen conditions. Based on our results, easy-to-hard curricula appear able to discover this low-dimensional, essential feature space (Figs 4D, S2D, S3F, S6B).

### Lower-dimensional readout subspaces contain more orientation selective neurons

Motivated by our hypothesis that models trained on easy-to-hard curricula appear to be relying on fewer essential visual features and are therefore more robust to transfer conditions, we further investigated the properties of the readout subspaces formed when training on each curriculum. To determine whether the readout subspaces are composed of similar features, we investigated whether the specific neurons which composed the learned readout subspaces were the same across curricula. We compared the indices of the 150 neurons in the readout subspace across curricula using the Jaccard index, which measures the similarity of two sets. A Jaccard index of 1 means the sets are identical, and a Jaccard index of 0 means the sets have no elements in common. We found that models trained on the shuffled and sequential curricula have the most readout neurons in common, followed by the shuffled and non-sequential curricula, while the sequential and non-sequential curricula share the fewest readout neurons (Fig 5A). To confirm that within-model variability does not underlie this difference across curricula, we also measured the Jaccard index across curricula normalized by the average Jaccard index within a single curriculum and observed the same similarity pattern (Fig 5B). These results support our claim that the curricula read out from different subspaces at the end of training. However, this also indicates some presence of shared features in the learned solutions of the shuffled and sequential models, given their high similarity. We next investigated these shared and unique visual features through orientation tuning analysis.

**Fig 5.**
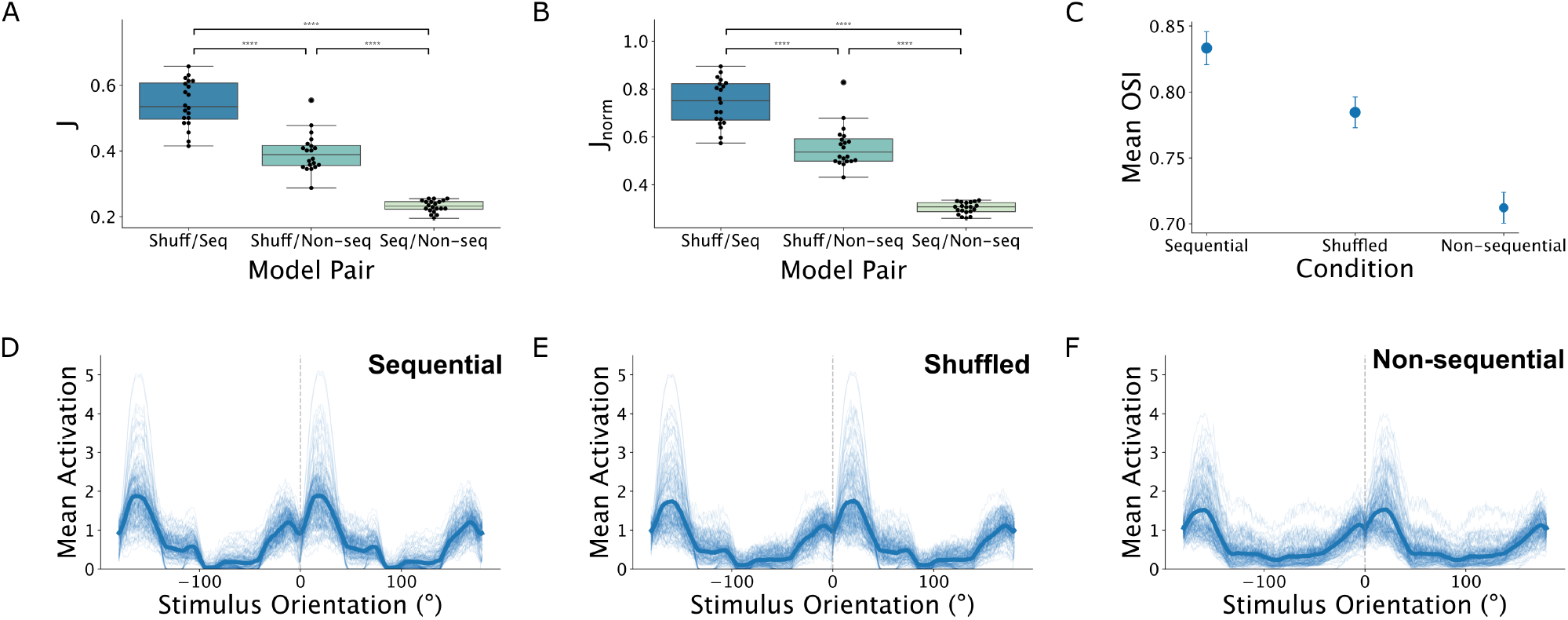
Lower-dimensional readout subspaces contain highly orientation-selective neurons. **(A)** Jaccard index for the set of indices of neurons in the readout subspace compared across curricula. Models trained on shuffled or sequential curricula have the most neurons in common. **(B)** Jaccard index for the set of indices of neurons in the readout subspace compared across curricula, normalized by the Jaccard index between models trained on the same curriculum. **(C)** Mean orientation selectivity of the 150 neurons in the readout subspace, measured by the orientation selectivity index, for sequential, shuffled, and non-sequential curricula. Curricula with a lower-dimensional readout subspace (Fig 4D) have a higher selectivity index. **(D)** Mean activations of each readout subspace neuron across stimulus orientations in the sequential curriculum. The mean across all neurons is shown as a thick line, individual neurons are shown as thin lines. **(E)** Same as (D) but in the shuffled curriculum. **(F)** Same as (D) but in the non-sequential curriculum. Both sequential and shuffled curricula have more consistent orientation selectivity across neurons than the non-sequential curriculum.

To solve the orientation discrimination task, during training, the models must identify and rely on neurons that are maximally informative regarding stimulus orientation. These orientation-selective units detect the essential visual features necessary and sufficient for task performance. To investigate whether the various models differ in their reliance on these units, we quantified the orientation selectivity of the most influential neurons within each model. Orientation tuning analysis on the influential neurons in the readout subspace showed that there was a difference between the tuning properties of the neurons depending on the curriculum. We measured the difference in tuning properties using the orientation selectivity index (OSI), which measures the selectivity of each neuron for its preferred orientation compared to the selectivity for the orthogonal orientation [37]. More selective neurons have a higher OSI. The neurons in the readout subspace of the sequential model were more selective for stimulus orientation than the other two models, and the neurons in the readout subspace of the shuffled model were more selective than the non-sequential model (Fig 5C). When we visualized the tuning curve of each neuron, we found that qualitatively, the neurons in the sequential and shuffled readout subspaces have more consistent orientation selectivity than the non-sequential curriculum (Fig 5D-F). This indicates that the sequential and shuffled curricula lead to low-dimensional readout subspaces which also rely mainly on stimulus orientation (i.e., more orientation selectivity), while the nonsequential curricula leads to a broader, high-dimensional set of readout neurons with less orientation selectivity. This supports our claim that lower-dimensional readout subspaces generalize better because they use fewer essential visual features and do not rely on non-essential visual features.

### Early training steps set the dimensionality of the readout subspace

Given our main theoretical claim that generalization depends on the dimensionality of the readout subspace, we examined the formation of readout subspaces across models throughout training. Throughout training, we measured the Jaccard index for the set of the indices of the top 150 influential neurons, in comparison to the set of the indices of the most influential neurons from the previous timestep (Fig 6A). We found that the Jaccard index plateaued at 1 in the first 200 steps of training, meaning the readout subspace was mainly set in the first 200 steps of training. There was a small change in the readout subspace for the sequential curriculum at steps 500-600, which is expected when the task condition changes from easy to hard and the model adapts to the previously unseen samples. This indicates that the initial steps of training mainly set the most influential neurons that form the readout subspace. Because the dimensionality of the readout subspace is determined by the neurons included in the subspace, this means that the initial steps of training mainly set the dimensionality of the readout subspace.

**Fig 6.**
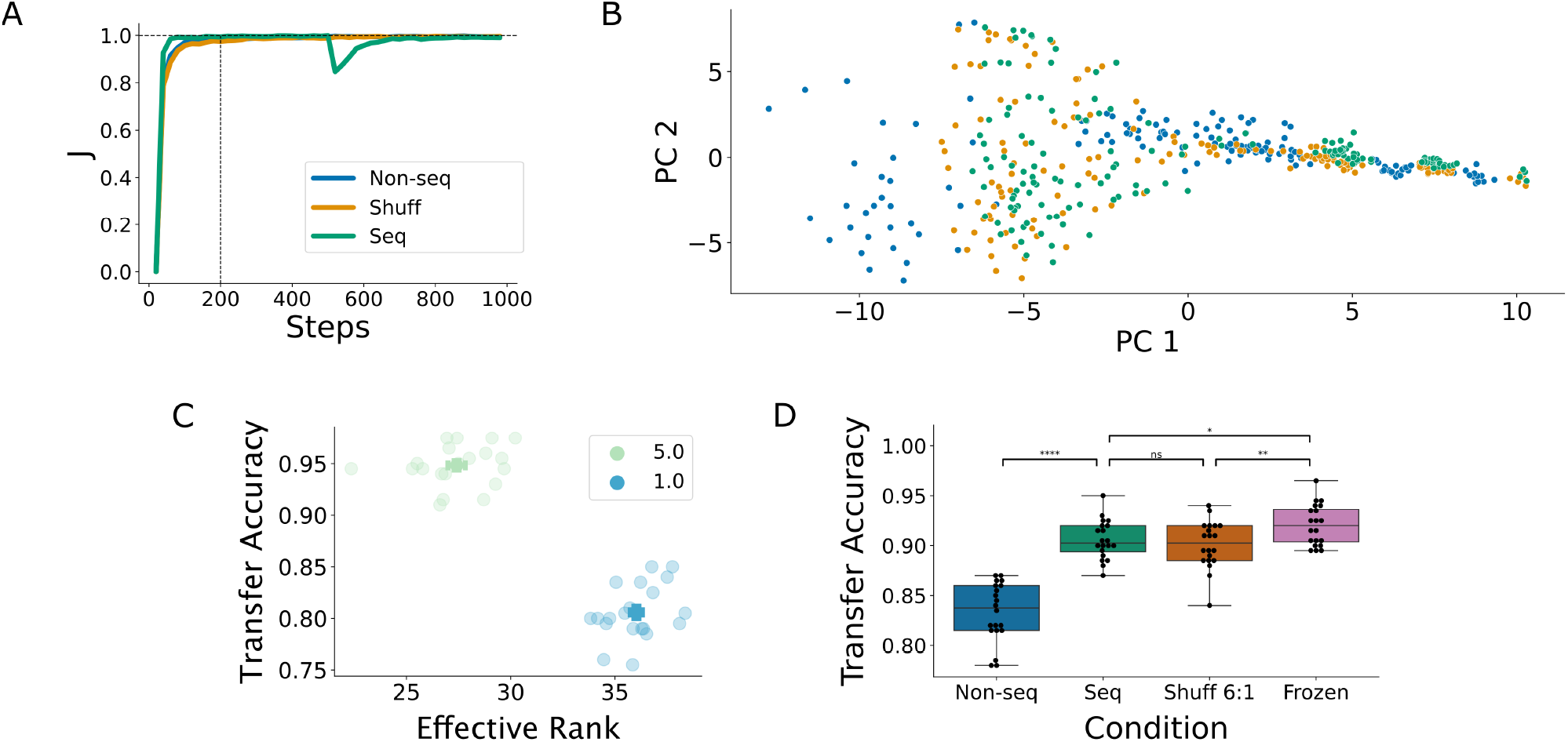
Readout subspace dimensionality is set early in training. (**A**) Jaccard index for the set of indices of the top 150 neurons, compared to the set of indices of the top 150 neurons from the previous step of training. The neurons involved in the readout subspace were set in the first 200 steps of training, and did not change much as training progressed. There was a small drop in the Jaccard index for the sequential curriculum at step 500, when the condition switched from easy to hard. (**B**) PCA of final readout subspace activations for a representative model. The activations in the readout subspaces form a continuum in PCA space from non-sequential → shuffled → sequential, with the shuffled and sequential curricula leading to very similar readout subspaces compared to the non-sequential curriculum. (**C**) Effective rank vs. task difficulty. Training on easy tasks led to a lower-dimensional readout subspace and greater generalization. (**D**) Comparison of a non-sequentially trained model constrained to use only the 150-neuron readout subspace transplanted from the forced sequential model. This allowed the model to learn and generalize on a hard task by searching within a pre-established low-dimensional subspace, improving generalization.

We also examined the representational properties of the neurons influencing the readout subspace and compared them across our three base curricula (non-sequential, sequential, and shuffled). To this end, we measured the activations of the 150 influential neurons in each model in response to 100 natural images (see Methods for further details), and after dimensionality reduction using principal component analysis, we compared the response profiles of the three sets of influential neurons (Fig 6B). The representational profiles of the neurons for the three curricula appear to vary on a continuum. Interestingly, the non-sequential subspace had the least overlap with the other two readout subspaces. The shuffled and sequential subspaces, however, were much more similar to each other in terms of their representational profiles, indicating that, as expected, training curricula that involve both easy and hard examples lead to more similar readout subspaces.

What causes the differing dimensionalities of the readout subspaces across curricula? By training solely on easy or hard tasks, we found that the easy task (i.e., larger angle separation) led to a lower effective rank of readout subspace than the hard task (i.e., small angle separation) (Fig 6C). Because the readout subspace is mainly set during the initial steps of training, and the following phases of training appear to stay close to the formed initial readout subspace, it follows that beginning training with an easy task helps to set a lower-dimensional, more generalizable subspace that is then reused for more difficult tasks. This provides a mechanistic explanation for why certain observers in the shuffled condition outperformed others: the observers who performed better focused on easy tasks first (Fig 3A), establishing a low-dimensional subspace that could be reused for the rest of training. Intuitively, the low-dimensional readout subspace discovered during the easy training phase proves sufficiently effective for the subsequent hard task that the training process preserves rather than substantially alters this configuration. This leads us to the final part of our main theoretical claim: less generalizable tasks can “piggyback” [7] on the low-dimensional subspace established through a more generalizable training task, leading to better generalization.

We directly tested this claim by training a model non-sequentially on the hard task but enforcing a learned low-dimensional readout subspace on the model. This mimics the readout subspace set by an easy task at the beginning of training. We enforced the low-dimensional subspace by freezing the readout weights of all neurons except for 150 neurons, the locations of which we adopted directly from the low-dimensional readout subspace of a shuffled model trained with a 6:1 easy:hard sample ratio. We chose this ratio because it led to the highest transfer accuracy (Fig 3E). This allowed the “frozen” model to learn and generalize on a hard task by searching within a pre-established readout subspace. Indeed, we found that limiting the learning subspace of the model to a lower dimension significantly improved generalization on the hard task (Fig 6D; *p*<<0.0001). We also found that the frozen model performed significantly better than both the sequential (*p*=0.025) and shuffled with 6:1 sample difficulty ratio (*p*=0.0084) models (Fig 6D). The superior performance of the frozen model compared to the model from which the readout subspace was derived likely reflects an additional regularization effect created by constraining learning to fewer neurons.

## Discussion

We compared ANN models to human observers on three different perceptual learning curricula: an easy-to-hard sequential curriculum, a shuffled curriculum with randomly interleaved easy and hard tasks, and a baseline non-sequential training paradigm. Human observers showed similar generalization improvements in both shuffled and sequential curricula compared to the non-sequential condition, while ANN models showed significantly greater generalization in the sequential curriculum than in the shuffled curriculum. We showed that both models and observers followed an implicit easy-to-hard curriculum, and that the strength of this curriculum correlated with generalization. To explain generalization differences across curricula, we proposed a theory based on readout subspace dimensionality. We showed that the dimensionality of readout subspaces (neurons with distinct visual feature preferences that contributed strongly to model behaviour) correlated with generalization capacity. Importantly, the readout subspace and its dimensionality were largely determined by task order during learning. Below we discuss the relevance of our findings in the context of previous research in perceptual learning as well as deep learning and artificial intelligence.

### Generalization in VPL

Generalization in perceptual learning has been investigated for decades [4]. As shown in previous works, many factors in tasks [38], paradigm design [8,39], and stimuli [6] influence generalization, but we yet lack a comprehensive theory of how all these factors together have an impact on learning generalization. Previous theories attributed learning generalization mainly to the receptive field properties of sensory neurons that were considered to contribute to the learned behaviour [30,40] or were re-tuned during learning [41]. Despite their success in explaining certain generalization variability, their explanatory power is mostly limited to the relationship between learning generalization and stimulus content. Other factors, such as learning sequence or curriculum, cannot be easily and directly explained by the existing theories. The problems of generalization and its underlying factors have also been a central research question in machine learning, and in particular, deep learning and ANNs [42]. Until recently [17], there has not been an explicit attempt to bridge between the two domains and connect the concepts of generalizations in machine learning and VPL, even though the observed phenomena are, at least from a behavioural standpoint, identical. Furthermore, potential empirical and theoretical contributions can easily be imagined in both directions. As a first step in this direction, as shown in this work and previous studies, we can start by investigating whether same patterns of generalization across training conditions and tasks emerge in ANNs and humans. For example, in the case of orientation discrimination and different angle separations, we observed that a similar pattern of generalization emerges in the two systems: an inverse relationship between learning generalization and angle separation is shared in the two systems [17]. We can then leverage the ANN models, and benefit from the accessibility to their whole inner machinery, to formulate a computational and mechanistic understanding of why different training paradigms, task designs, stimuli, or curricula lead to different generalization abilities. Theoretical studies of generalization in ANNs have shown fundamental advances in the last few years [42–45] that could offer new insights into perceptual learning generalization in humans. Equipped with new theoretical ideas, we can then develop new experiments that test neuronal and behavioural predictions following from this computational understanding.

Following this approach, we propose a theory of perceptual learning generalization: different training tasks, stimuli, and curricula lead to readout from different sensory populations with varying dimensionalities. As we demonstrated in our ANN model, this readout dimensionality strongly predicts learning generalization. Importantly, this theory has specific experimental predictions for how sensory representations in the brain are communicated to downstream sensorimotor or association areas. Specifically, the dimensionality of the communication subspace between sensory areas and downstream areas should depend on the training task, and we should be able to predict the learning generalization by measuring this dimensionality of the communication subspace. Although inter-area communication subspaces have been studied recently [46– 48], the link between them, prior learning experience, and the characteristics of learned behaviour requires further investigation. The main challenge lies in investigating the communication subspace between sensory and sensorimotor areas that are causally involved in the learned task. Although experimentally challenging, such investigation is feasible [26,49]. This clear experimental prediction provides opportunities for further validation and refinement of our theory, which can bridge behaviour and its underlying neuronal mechanisms.

### Learning curriculum

In deep learning literature, there are two views of the utility of learning curriculum. The first claims that easy-to-hard, or simple-to-complex, ordering of training data can help models learn faster and generalize better [33,50–53]. The other, more recent, view claims that in training regimes with large datasets and large models, the explicit ordering of the training samples does not play a significant role in generalization, and it does not benefit models to start with easier examples [54,55]. However, in training regimes with small datasets, sparse [56] or noisy [31,57] data, small models, or short training times [31], curriculum learning may still play a significant role. In particular, a curriculum can increase the efficiency of training, requiring smaller datasets and shorter training times [58–60]. Interestingly, as shown recently, learning curriculum may help models behave more similarly to biological organisms [61].

VPL is generally studied as a training regime with small training datasets and short training times, particularly when compared to the training regimes of large ANNs. If the same principle applies to human learning, perceptual learning should improve with curriculum learning. Our results indeed corroborate this expectation: a sequential learning curriculum improved learning speed and generalization. However, once human observers were offered an unstructured training curriculum with a shuffled order of task difficulty, they proved to be capable of following more complex learning strategies that were broadly as effective as an explicit sequential learning curriculum. The model, on the other hand, was unable to discover these more effective learning strategies from an unstructured training curriculum. As a future direction, it will be important to quantify curriculum efficacy as a function of training stimulus set size in humans. If the similarity with ANNs holds, we expect to see reduced curriculum effects as training stimulus sets increase in size.

In machine learning, previous work focusing on “active learning” has developed both heuristics and principled algorithms [62] for choosing an order for training examples to learn from. We find that our model benefited from an explicit sequential or quasi-sequential ordering, consistent with previous works that showed the benefits of an easy-to-hard curriculum for ANNs in limited training time and samples [31,56,58–60].

### Inter-individual variability in learning

Inter-individual variability has not generally been emphasized in VPL, and common practice is to average over a large group of individual learning trajectories to discover shared learning strategies [63]. However, our results suggest that inter-individual differences may play a large role in visual learning. Our analysis of human observers suggests that there is substantial variability in the implicit curriculum learned from tasks with randomly interleaved difficulty levels. In this paper, we found that this variability led to different learning outcomes, and that the strength of the implicit curriculum predicted generalization (Fig 3C-E). This interpretation could help to reconcile the differing conclusions of previous literature on the benefits of a sequential curriculum [6–9]. Averaging across participants using an interleaved curriculum removes the inter-individual variability, obscuring the conclusion that some participants may benefit from a shuffled curriculum while some benefit from an explicit sequential curriculum. Consistent with our results, recent work has shown that there are substantial inter-individual differences when learning visual tasks [63–65]. As shown in [65], mice displayed notable inter-individual differences in learning strategy and trajectory throughout training, and the strategy used later in training was predictable from the strategy used at the beginning of training.

It is important to note that, while we have studied inter-individual variability in the context of behavioural responses and generalization performance, there are likely multiple underlying factors at play. Previous work by Yang et. al focused on the baseline abilities of individuals to learn a perceptual task, which they hypothesized could underly individual differences in perceptual learning [63]. The authors were able to extract a subject-specific and task-invariant “perceptual learning ability” which measured an individual’s general baseline ability to learn the task, and which comprised a large part of the overall variance in learning [63]. This suggests that individual differences in learning strategy or ability may drive performance on perceptual learning tasks, consistent with our findings that the observers’ implicit curricula were a main driver of differences in generalization performance in the shuffled condition. In addition to varying behavioural responses and perceptual abilities, the computations performed by neurons in response to visual stimuli and during deliberation, which then lead to behavioural responses, also vary across individuals [66]. These varying neural computations could be responsible for the wide variability both in baseline perceptual learning ability [63] and in learning strategy [64,65,67] observed in our study and in previous work, as different neural computations lead to different behavioural responses and therefore may underly differing performance and generalization abilities.

### Learning and reusing low-dimensional subspaces

In this paper, we hypothesized that learning from easy tasks early in training forms a low-dimensional subspace in which fewer independent visual features contribute to visual decisions. We showed that reuse of this low-dimensional subspace drives generalization to harder tasks learned later in training. This theory shares commonalities with previous work on visual learning in humans and ANNs [43,68,69], which converges on a similar theoretical framework from different perspectives.

Previous work in deep learning has provided a theoretical explanation for why different training tasks may lead to different generalization [43]. The spectral theory of learning generalization claims that the spectral properties of a representational space – namely, the distribution of eigenvalues and eigenvectors of neural responses to stimuli [68] – and how the training task (i.e., input-output mapping) aligns with the population’s eigenspectrum determines the resulting generalization. More specifically, tasks that align with the network’s top eigenvectors and that can be solved using these low-dimensional eigenvectors lead to better generalization in few-shot learning regimes. In our experiments, easier tasks (e.g., larger angle separation in orientation discrimination) may be better aligned with the top eigenvectors of the underlying population, hence are learned faster and generalize better. This is consistent with our observation showing that a larger angle separation led to a lower-dimensional readout (Fig 6C), representing the complementary perspective of the spectral theory. Prior experimental evidence in animal studies also supports the possibility of low-dimensional readouts under certain training conditions [49,70,71].

Beyond what was proposed by the spectral theory of learning generalization, we also showed that a hard training phase (low alignment with top eigenvectors, using the terms from [43]) that follows an easy training condition can leverage the same recruited low-dimensional subspace as the easy task and achieve better generalization compared to training only on the hard task. This is consistent with previous empirical observations in humans showing that a second training task can “piggyback” on the generalization outcome of a first training task [7].

Extending beyond visual learning, this theory is also consistent with work in dynamical systems which found that neuronal motifs (i.e., low-dimensional neural representations of task structure) are reconfigured and reused across similar tasks [72]. Previous work has shown that forcing a quasi-sequential implicit curriculum through gradual learning from a “teacher” model improves learning speed, generalization, and overall performance [73]. Learning from a teacher model in this way gradually scales up the complexity of the task and input representations, thereby enforcing a low-dimensional representation for the “student” model at the beginning of training that can be reused for more complex tasks [73]. This reinforces the benefits of our “forced” quasi-sequential curriculum in aiding the model to learn and reuse a low-dimensional readout subspace which it would not otherwise discover on its own.

### Applications to real-world curricula

While the scope of this study is restricted to simple visual learning tasks, there are potential applications of our work in developing real-world curricula for perceptual generalization. For example, medical fields where diagnoses are made visually require a large amount of perceptual expertise and the ability to generalize to novel abnormal images across individuals [74,75]. For this reason, training the perceptual abilities of radiologists or other specialists who rely on their perceptual insights for diagnosis is extremely important. Although there is no generally accepted perceptual learning curriculum for these fields, it has been shown that initially training students on easier examples before progressing to complex examples can lead to significant improvements in recognition of abnormal images [76,77].

Our findings can also extend beyond the purely perceptual realm. Studies on classroom curricula have found advantages in both easy-to-hard sequential training [78] as well as interleaving various aspects of learning [79–81]. In our results, we found no significant difference between interleaving, or shuffling, tasks vs. ordering them sequentially in an easy-to-hard manner. However, we also found that some observers can benefit from the shuffled curriculum, as it allows them to develop their own implicit sequential curriculum near the beginning of training that ultimately leads to better generalization than an explicit sequential curriculum. This suggests that in a classroom setting, interleaved curricula may be useful to allow students to develop their own individual implicit curricula. However, as shown in our results, not everyone will benefit as strongly from interleaved learning, and might be better suited to an explicit sequential curriculum. For example, an explicit sequential curriculum has been suggested for students with learning disabilities [78]. Therefore, the optimal curriculum for learning generalization may vary across individuals as well as across tasks.

Additionally, most education involves multiple sensory modalities and task types. Therefore, the optimal curriculum for learning classroom material may be much more complex than the optimal curriculum for visual learning generalization. Supporting this, studies have shown that more complex curricula [79,80,82] that may involve both sequential and interleaved components [83,84] are optimal for different tasks in both humans and ANNs. For example, recent work in ANNs has suggested that in both humans and ANNs, interleaving is advantageous during incremental learning and blocking is advantageous during rapid inference [84]. On the other hand, a recent study comparing transformers and ANNs on a rule-learning query response task found that humans benefit from observing many diverse examples at the beginning of training to promote in-context learning first, while ANNs do not benefit [85]. These studies point towards the need for a more complex curriculum as the complexity of the subject matter increases. Future work could build on our results and theory of generalization to develop and compare these optimal curricula across tasks and perceptual modalities, in addition to across individuals.

### Limitations

One limitation of our study is the use of backpropagation as the learning algorithm. Backpropagation, which calculates loss gradients with respect to model parameters in the gradient descent algorithm used in virtually all machine learning models, including for modeling VPL [17], has known issues with biological plausibility [86,87]. Additionally, gradient descent learned through backpropagation violates Dale’s Law and produces biologically implausible weight distributions [88]. Therefore, the fact that our model did not closely follow the same implicit curriculum as human observers in the shuffled condition could be due or partially due to our use of backpropagation. To address this limitation, future work could reproduce our experiments using biologically plausible learning rules [89].

One other limitation of our analysis is our choice of curriculum metric. In this work, we used the difference between the mean accuracy on the first *n* easy trials and the mean accuracy on the first *n* hard trials as a proxy for the observers’ implicit curricula. While this metric isn’t a direct measure of curriculum, we chose it as a proxy in order to deal with the inherent noise present in the observer accuracy data. A rate of change metric would better approximate the actual implicit curriculum, as would a trial-by-trial metric instead of averaging over *n* trials, but due to the noisiness of the data these are technically infeasible. In future work, more advanced techniques for measuring the strength of the implicit curriculum could be effective.

A final limitation of our work is the pre-trained ANN architecture that we used in our modeling experiments. AlexNet [21] has often been used to model the ventral visual pathway in previous work on VPL [17,90,91] and has been shown to perform similarly to newer and more complex architectures in terms of model-brain alignment [24]. Further justification of our choice of model was presented in the results section. However, AlexNet and similar architectures remain rough approximations of visual system architecture and connectivity. For example, feedforward networks like AlexNet lack the lateral or feedback recurrent connections present in biological systems (91). Some have argued that models combining feedforward and recurrent connections are more plausible models of VPL than AlexNet or other strictly feedforward ANNs (91, 92). In an effort to address this limitation, we investigated several other model backbones (see S1 Appendix, S7 Fig). Although the correlation between readout subspace dimensionality and generalization holds for some models, it is not universal across all backbone architectures. This variability indicates that the relationship between readout dimensionality and generalization may interact with specific architectural properties. Further research is required to characterize how different architectural inductive biases influence the emergence of this dimensionality-generalization relationship.

## Materials & Methods

### Ethics Statement

Observers gave written, informed consent before their participation, and the study was approved by the Ethics Committee of the Université de Montréal (CEREP, Project no. 2024-5806).

### ANN Model

#### Stimuli

Gabor stimuli for training the ANN models were generated through the PsychoPy software [92], following the parameters described in Wenliang and Seitz [17]. Briefly, each image was 227×227 pixels, centered on a gray background. A reference stimulus with 0° orientation was used in all experiments. Various target stimuli were used as described in the results section, with angle separations between the reference and target stimulus ranging from 0.5°-10.0°. We used a spatial frequency (SF) of 0.05 (spatial wavelength (SW) = 20px) for the train condition and a SF of 0.1 (SW = 10px) for the transfer condition. We added a Gaussian mask with a SD of 50 pixels to each image. For the additional experiments in S2 Fig, we used a reference stimulus with 0° orientation for the train condition and a reference stimulus with 15° orientation for the transfer condition, with a constant SF of 0.05.

#### Architecture

We simulated visual learning using an ANN based on AlexNet [21], with modifications based on previous work by Wenliang and Seitz [17] in addition to our own modifications. Briefly, our model has five convolutional layers, three max-pooling layers, and one fully connected readout layer (Fig 1A). We modified the model from the version proposed by Wenliang and Seitz [17] to add skip connections from every layer of the model directly to the readout layer. The weights in the readout layer were initialized to zero to simulate the instruction period of the task [17], and the weights for all other layers were loaded in PyTorch from an AlexNet model trained on object recognition on the ImageNet dataset [22]. To confirm that zero initialization does not affect the main conclusions, we repeated the main experiments with a randomly initialized readout weight vector (S3 Fig). For the results in S7 Fig, we substituted other model backbones for the AlexNet architecture and added skip connections from every layer of the model directly to a single readout layer. We initialized the readout layers randomly for these experiments and froze the model backbones, allowing only the readout layer to update. The results shown in S7 Fig include the AlexNet, EfficientNet, and GoogLeNet backbones. We also tried more complex architectures: ConvNeXt, DenseNet121, MNASNet, MobileNetV1/V2, SqueezeNet, ResNet18, and VGG11/16. See S1 Appendix for details.

#### Training

To simulate human error due to both noise in viewing the stimulus and uncertainty about the decision, we added Gaussian noise to both the stimuli (SD = 5px) and the confidence estimate of each model prediction (SD=0.3) at runtime. We applied this noise independently for each trial. We used single sample updates (i.e. batch size of 1) to train the model to match the human observer training [64], as the observers only saw one sample per trial. For the results in S7 Fig, we used a batch size of 20 because some of the model backbones contained BatchNorm layers. We trained the model using the binary cross entropy loss function and stochastic gradient descent with backpropagation in PyTorch (learning rate = 0.0001) [17]. For the results in Figs 1, S1, S2A-B, and S3A-D, we trained 100 models each on a single angle separation (0.5°, 1°, 2°, 5°, or 10°) for 500 iterations each and tested the transfer to the same angle separation. For the results in Figs 6C and S7, we trained 20 models each on a single angle separation for 1000 iterations and tested the transfer to the same angle separation. For all three curricula (non-sequential, shuffled, and sequential) we tested the transfer on a 1° angle separation (i.e., the hard condition). For the results in Fig S2C-E, we trained 30 models on each curriculum for a total of 1000 iterations due to the increased variability between curricula. For the results in Fig 4C, we trained 100 models on each curriculum due to high variability in the lesioning process. For all other results, we trained 20 models on each curriculum for a total of 1000 iterations each as outlined below.

### Experimental design

#### Observers

37 total observers with normal or corrected-to-normal vision participated in the experiment (14 male observers, 23 female observers; age, 22.11±3.58 years; range, 16-29 years). The experiment was run remotely and was controlled by a PsychoPy [92] program that observers downloaded and ran on their own computer. Participants completed the study at home, on personal computers. Stimulus parameters were calibrated to each observer’s screen size and their reported viewing distance. Participants were asked to choose a viewing distance within a range of 35-60cm from the screen and sit at the same distance from the screen on each day of the experiment.

#### Stimuli

Gabor stimuli for the human observer experiments were also generated through PsychoPy, following previous work by Jeter et al. [18]. Stimuli subtended 3° × 3° visual angle at a viewing distance of 35-60cm. The observers were allowed to select a viewing distance within this range but were instructed to keep the viewing distance consistent across days. The sizes of the stimuli were calculated based on the viewing distance and the screen size of the device used for the experiment. Stimuli were presented in the periphery, 3° out from the centre of the screen. Stimuli were randomly presented on either the upper-left or bottom-right corner of the screen with a reference angle of 145° during training and presented on either the upper-right or bottom-left corner with a reference angle of 55° during transfer. Target stimuli were rotated either 5° or 1° clockwise or counterclockwise from the reference orientation, according to the training paradigm. Stimuli were presented on a gray background.

#### Training

On each trial, observers fixated on a small bullseye fixation point in the centre of the screen for 280±20ms. A reference image was then shown for 300ms, followed by a 280±20ms break, and then a target image for 300ms. After 750±20ms, observers were asked to respond with a keypress to indicate whether the target stimulus was rotated clockwise (j key) or counterclockwise (f key) from the reference. The response time was capped at 10s, and trials where observers reached the maximum response time without responding were recorded as an incorrect response. Responses with a RT of ≥10s were discarded as non-responses for all calculations and plots in the results. During training, a green circle was briefly overlaid on the central fixation point following each correct response and a red circle was overlaid following each error. This visual feedback was removed during transfer. The next trial began 750ms after the keypress response.

We trained observers for one session per day on four consecutive days and tested them on the transfer condition for one session on the fifth day. During each session, observers were trained on four blocks of 80 trials each (1280 total trials). Observers were also tested on four blocks of 80 trials each during the transfer day. Each block was separated by a rest period of at least one minute, which the observer could decide to prolong at their discretion. Accuracy, response times, and the angle separation for the viewed stimuli were recorded for each trial. If the observers did not respond in 10s, the response was recorded as incorrect, and the next trial began.

We randomly split the observers into the three different curricula outlined below. For the results reported in Figs 2 and 6 observers were assigned to the non-sequential curriculum, 6 observers were assigned to the sequential curriculum, and 10 observers were assigned to the shuffled curriculum. For the results reported in Fig 3, we recruited 15 additional participants who were assigned to the shuffled curriculum to better investigate individual variation in that condition.

### Curricula

We trained both models and observers on three curricula: a baseline non-sequential curriculum, an easy-to-hard sequential curriculum, and a shuffled curriculum where easy and hard conditions were randomly interleaved. In the non-sequential curriculum, models and observers were trained on only the 1° condition throughout training. In the sequential curriculum, models and observers were trained on the 5° condition for the first half of training (500 trials for the models, 2 days or 640 trials for the observers) and then trained on the 1° condition for the second half of training for an equal number of trials. In the shuffled curriculum, models and observers were trained on randomly interleaved 5° and 1° trials. For the shuffled condition, the order of the 5° and 1° trials was randomized for each participant/model instance.

### Analysis

#### Statistical analysis

Statistical analysis for all results was performed using Seaborn, Scikit-Learn, and Matplotlib. All error bars for plotting were calculated based on the standard error of the mean unless otherwise specified. For all boxplots, boxes represent the inter-quantile range (IQR) and whiskers are drawn to 1.5*IQR with Seaborn. Data was tested for normality with the Shapiro-Wilks test, and statistical significance was computed using the Welch’s t-test. For all boxplots, the annotations used are: ns = not statistically significant (p>0.05); * = p≤0.05; ** = p≤0.01; *** = p≤0.001; and **** = p≤0.0001. Correlation coefficients and p-values for linear correlations were calculated using Pearson correlation and a two-tailed t test. Correlations for Fig 3C-D, 4D-E, S2D, S3F, S5, and S6B were calculated over every individual datapoint. The correlation for S7 Fig was calculated using the means across all model backbones for a single angle separation. Correlations for all other plots were calculated over the means across each training condition. 95% confidence intervals were computed using the standard Seaborn regplot implementation of the t-distribution, based on the standard error of the mean across observers.

#### Accuracy calculations

We calculated the transfer accuracy of each model as the average accuracy over 200 trials of the transfer condition on the 1° angle separation. For the observers, we calculated transfer accuracy as the average accuracy over all four blocks of day 5 (320 trials in total).

#### Implicit curricula calculations

We split the response time and accuracy for each observer by the difficulty of the trial (i.e. an angle separation of 5° or 1°). In Fig 3A, accuracy is averaged across each block of training (80 trials).

Our curriculum metric for each observer (Fig 3C and 3D) was calculated as follows:

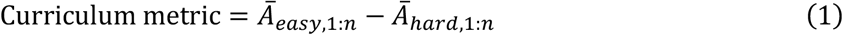

where Ā _*easy*,1: *n*_ represents the mean accuracy on the first *n* easy trials and Ā _*easy*,1: *n*_represents the mean accuracy on the first *n* hard trials. We chose to use this metric instead of, e.g., a rate of change metric, because calculating the difference in rate of change between consecutive trials considerably increased the amount of high-frequency noise in the data. Due to the high amount of noise between individual trials, we computed the metric over the first *n* easy/hard trials instead of computing the metric trial-by-trial.

To calculate the learning signal in the model, the mean was taken over all the absolute gradient values of the model’s training loss with respect to the model weights for the full model. The gradients were split by the difficulty of the trial. For Fig 3B, because only one difficulty condition occurs on each step of training, gradient magnitudes of the other condition were set to the previous value of that condition, or to zero if no previous value existed. For example, on a 1° trial, the gradient magnitude for plotting the 5° condition was set to the value calculated in the last 5° trial. For S4A Fig, we plotted both the active and non-active gradients at each timestep.

To approximate the range of implicit curricula found in human observers, we manipulated the ratio of sample difficulties while training the model on a shuffled curriculum (Fig 3E). We repeated the shuffled training as described previously, but instead of randomly selecting the task difficulty at each step with a 1:1 probability ratio of easy:hard samples, we modified this ratio to have a higher or lower probability of choosing each task difficulty. We used the modified ratio to choose samples for the first 25 steps of training and then returned to randomly sampling the task difficulties. We repeated the experiment with ratios ranging from 1:10 to 10:1 easy:hard samples.

#### Dimensionality analysis

For Fig 4A, PCA was calculated on the full readout weight vectors taken at the beginning of training, halfway through training, and at the end of training. For Fig 6B, PCA was calculated on the final activations of the 150-neuron readout subspace for representative models from each curriculum. The weights or activations from each model’s readout subspace were concatenated before PCA and then coloured by the model they originated from after PCA. For both PCA figures, we used the StandardScaler from SciKit-Learn to standardize the data to zero mean and unit variance before computing the top two principal components.

For Fig 4C, neurons were lesioned incrementally 10 at a time by setting their readout weights to zero. This removes the connection from the neuron to the readout. Lesioning was based on the absolute readout weight value, i.e. first the largest 10 readout weights were lesioned, then the largest 20 readout weights, etc. Transfer accuracy was measured as usual after each set of neurons were lesioned. The transfer accuracy was divided by the number of neurons lesioned to calculate the accuracy drop per neuron. To determine the functional contribution of individual neurons, every weight in the readout was lesioned one at a time and the transfer accuracy was measured as usual. After determining the 150 neurons that contributed the most to the readout, S6A Fig was created as described for Fig 4C.

To calculate the dimensionality of the readout subspace, activations were first collected [36] from the readout layers of the models before finetuning while passing 100 natural images from a subset of the ImageNet dataset [22] through the model. We selected 1000 images from the validation set of ILSVRC 2012 [93] as our subset. We selected 150 neurons from the readout of each model instance with the highest absolute weight values (or highest functional contribution as described above) and selected the activations from the models based on the indices of these neurons. This yielded a 100×150 activation matrix. We then performed SVD on each activation matrix and calculated the effective rank according to Roy and Vetterli [35].

The Jaccard index shown in Fig 5A was calculated as follows for each pair of models, for each trial *i*:

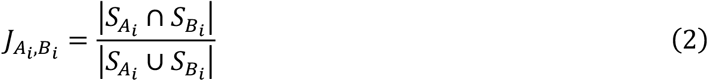

where A, B are models trained with different curricula, and 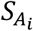 is the set of indices of the 150 neurons in the readout subspace of model A at the end of training for trial *i*.

In Fig 5B, the Jaccard index was normalized using the within-model variance:

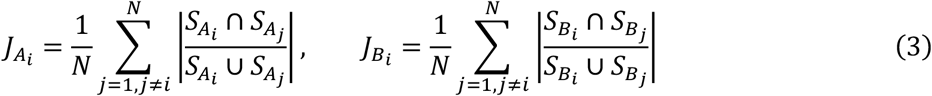

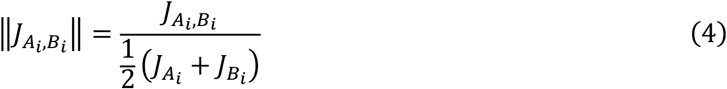

The Jaccard index shown in Fig 6A was calculated as follows for each model:

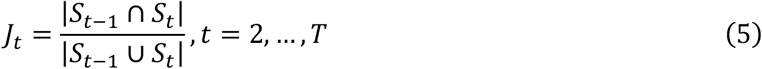

where *S*_*t*_ is the indices of the 150 neurons in the readout subspace at training step *t*.

#### Orientation tuning analysis

The orientation selectivity index (OSI) shown in Fig 5C was calculated using the following formula for each neuron in the 150-neuron readout subspace:

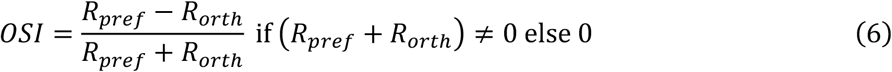

To calculate *R*_*pref*_, which is the firing rate, or activation, at the preferred orientation of each neuron, we first collected the activations from the readout layers of the models before finetuning while passing Gabor stimuli through the model. The Gabor stimuli had orientations of [-180, 180], and each was repeated 20 times. *R*_*pref*_ was then the mean maximum activation for that neuron (i.e., the maximum of the mean tuning curve). Then, *R*_*orth*_ is the mean activation of the neuron at the orthogonal orientation (preferred orientation + 90°). To plot the tuning curves shown in Fig 5D-F, we plotted the mean activation from each neuron for all orientations, and the mean across all neurons for all orientations.

## Acknowledgements

This work was supported by NSERC Discovery Grants (RGPIN-2023-03875 to SB; RGPIN-2023-03853 to CCP) and an IVADO Exploratory Project grant to SB and CCP (Explo24CO-3750823649). CV was supported in part by NSERC Undergraduate Student Research Awards. This research was enabled in part by support provided by (Calcul Québec) (https://www.calculquebec.ca/en/) and the Digital Research Alliance of Canada (https://alliancecan.ca/en). The funders had no role in study design, data collection and analysis, decision to publish, or preparation of the manuscript.

## Author contributions

Conceptualization: CV, CCP, SB

Methodology: CV, CCP, SB

Formal analysis: CV, SB

Writing - original draft: CV

Writing: Review & editing: CV, CCP, SB

Visualization: CV

Supervision: SB, CCP

Funding acquisition: SB, CCP

## Data and materials availability

All data is available in the main text, supplementary materials, and/or the locations specified in this section. The raw data from the human observers is publicly available at https://doi.org/10.6084/m9.figshare.32897081. All code necessary to reproduce the results is publicly available at https://github.com/charlottevolk/Curriculum-VPL.

## Supplementary Materials

**S1 Fig.**
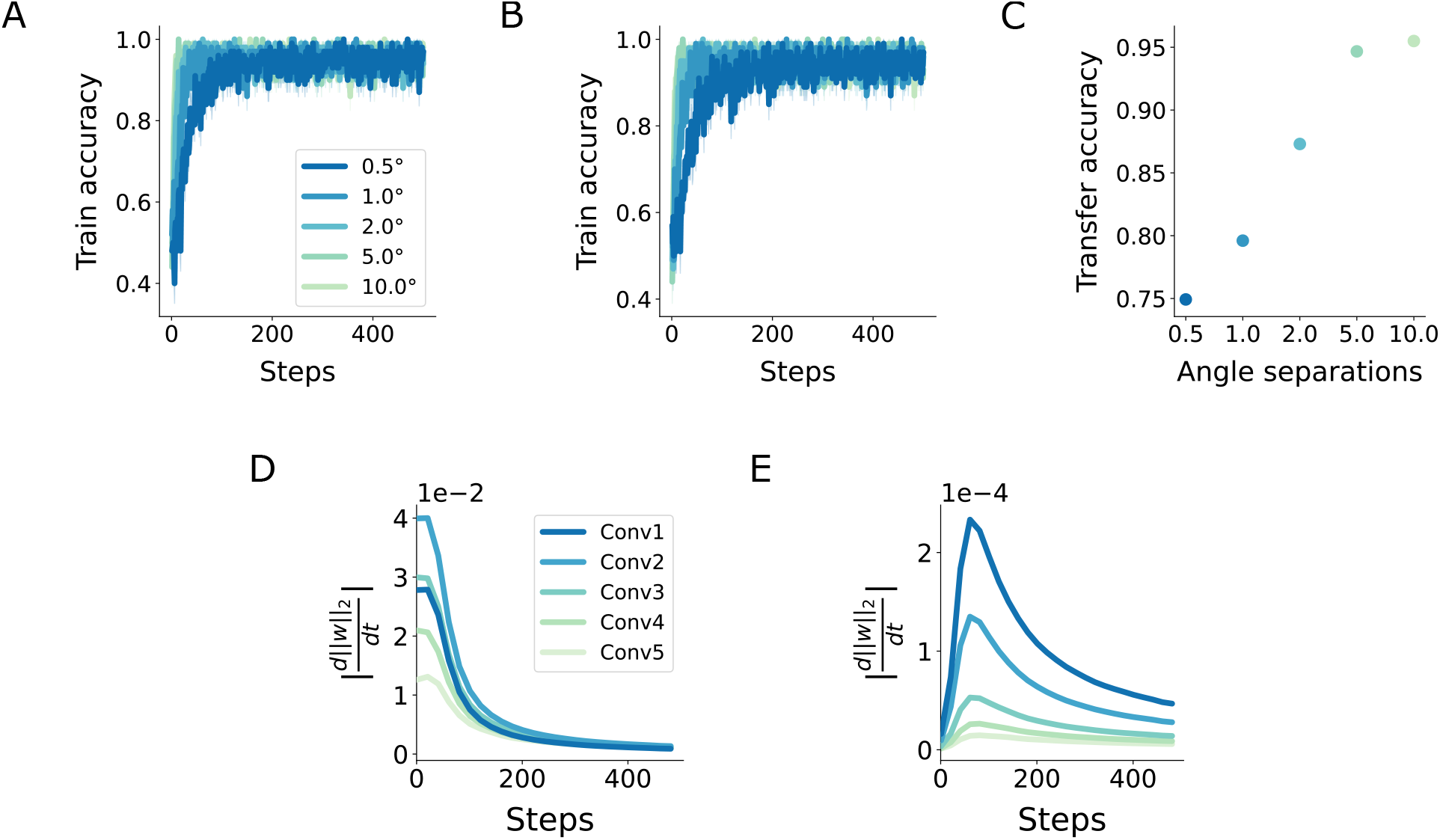
Additional training, transfer accuracy, and weight change data. (**A**) Train accuracy with standard error visible. (**B**) Train accuracy with standard error visible for a model with convolutional and pooling layer weights frozen and readout weights unfrozen. (**C**) Transfer accuracy for the model from B. **(D)** Gradient of the weight norm of skip connection weights throughout training on a 0.5° angle separation. **(E)** Gradient of the weight norm of convolutional weights throughout training on a 0.5° angle separation.

**S2 Fig.**
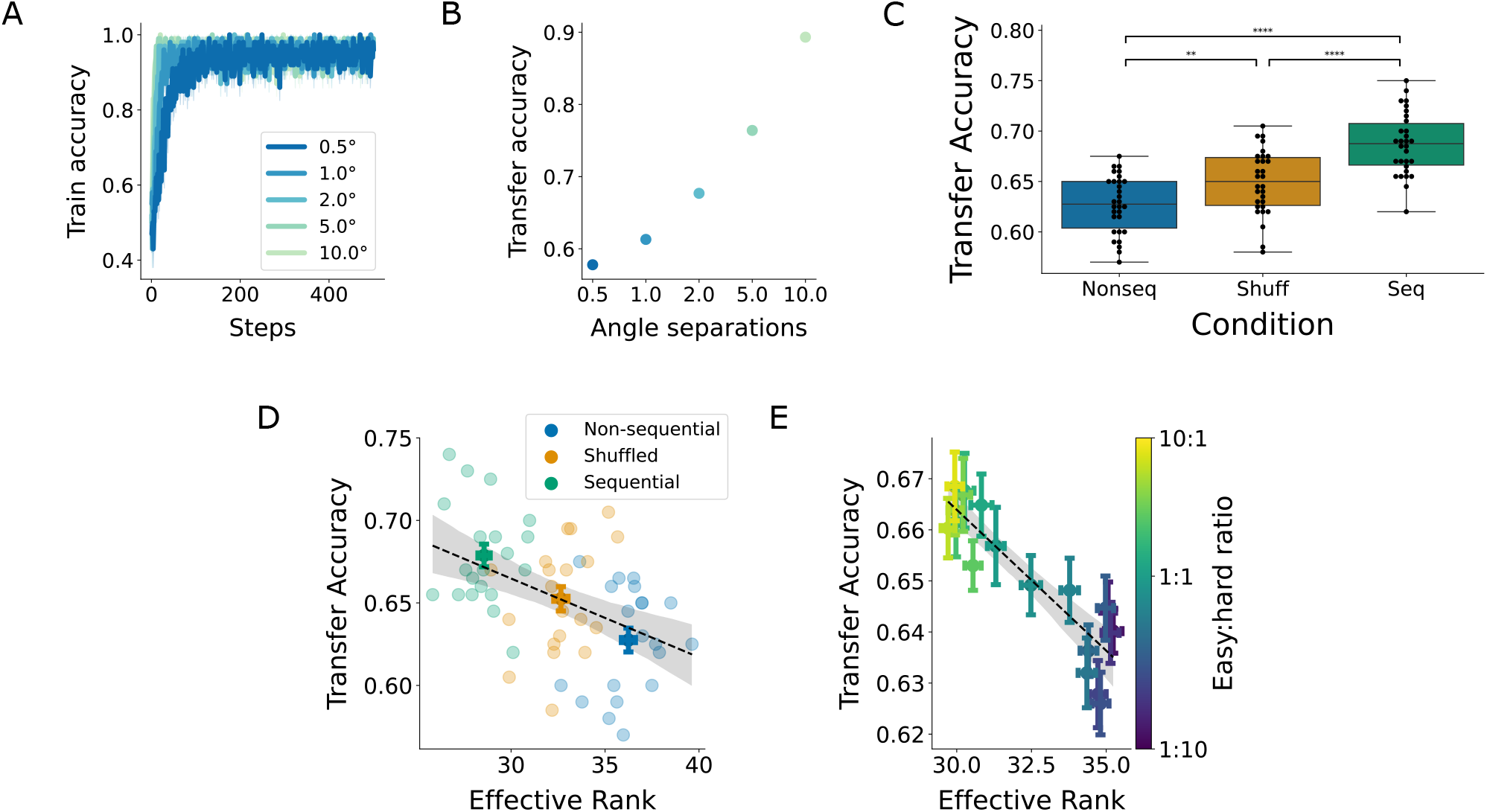
Results for a changed reference orientation in the transfer condition. **(A)** Train accuracy. **(B)** Transfer accuracy. **(C)** Transfer accuracy for models across curricula. **(D)** Transfer accuracy versus effective rank for models across curricula (*r*=-0.45, *p*=0.00034). **(E)** Negative correlation of the effective rank for shuffled curricula with different easy:hard sample ratios with transfer accuracy (*r*=-0.89, *p*<<0.0001).

**S3 Fig.**
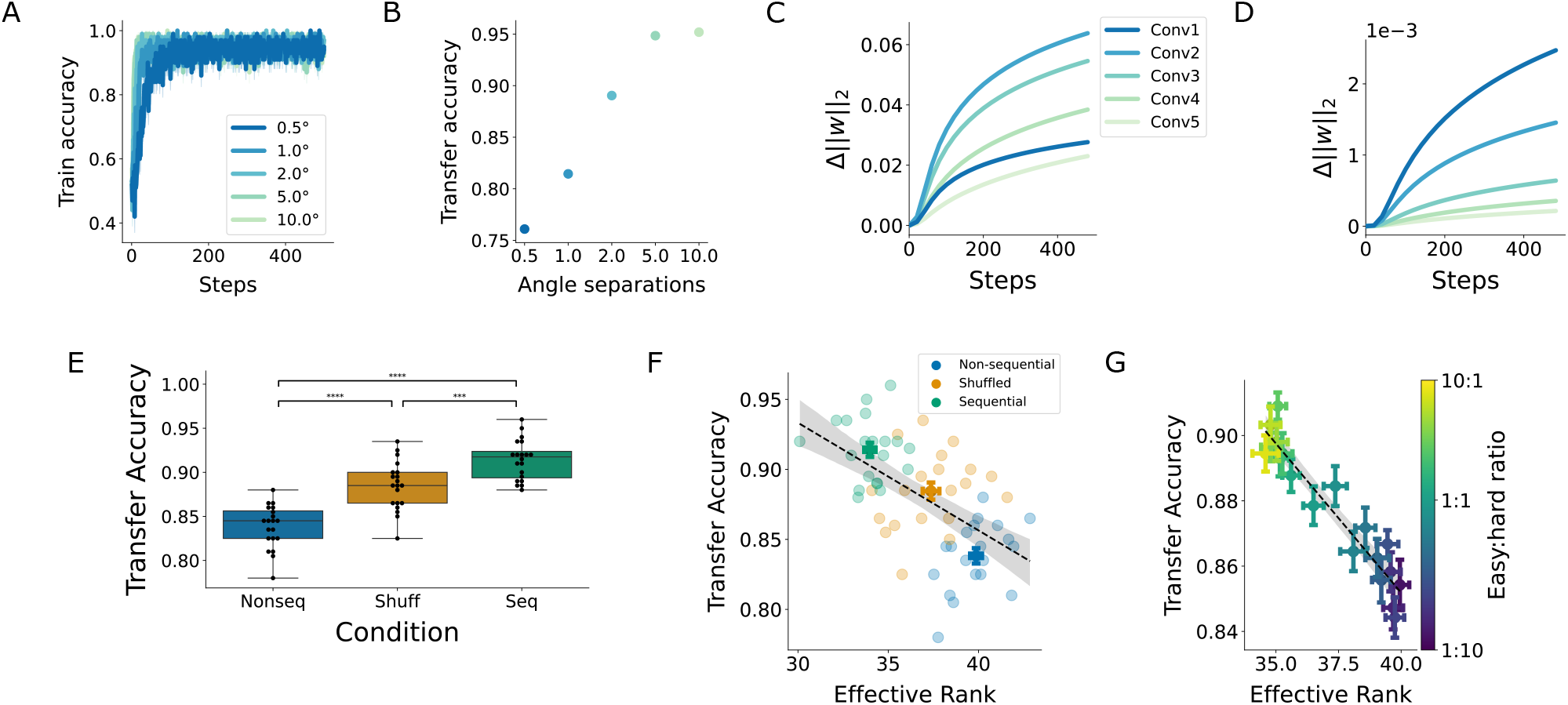
Results for models with randomly initialized readout weights. **(A)** Train accuracy. **(B)** Transfer accuracy. **(C)** Change in the L2-norm of skip connection weights throughout training on a 0.5° angle separation. **(D)** Change in the L2-norm of convolutional weights throughout training on a 0.5° angle separation. **(E)** Transfer accuracy for models across curricula. **(F)** Transfer accuracy versus effective rank for models across curricula (*r*=-0.57, *p*<<0.0001). **(G)** Negative correlation of the effective rank for shuffled curricula with different easy:hard sample ratios with transfer accuracy (*r*=-0.95, *p*<<0.0001).

**S4 Fig.**
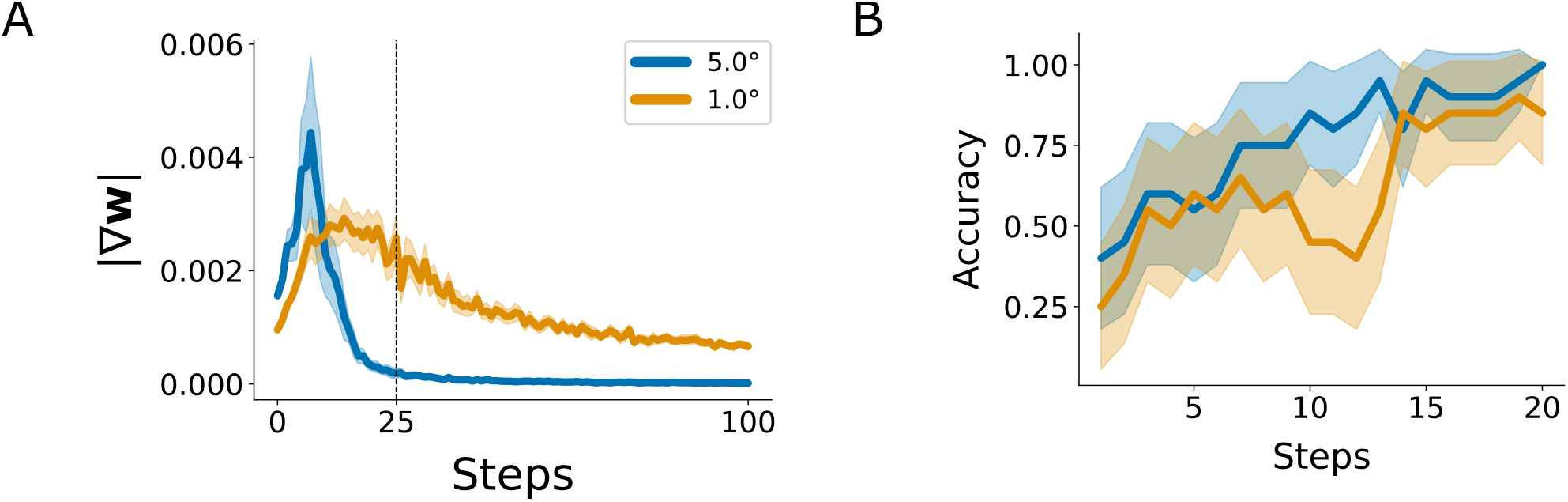
Additional data supporting an implicit curriculum in the models. **(A)** Parameter gradients for models trained on the shuffled task condition in the first 100 steps of training. This plot includes gradients for the non-active condition at each timestep. **(B)** Accuracy throughout training for models on the shuffled condition shown separately for easy (5°) and hard (1°) samples.

**S5 Fig.**
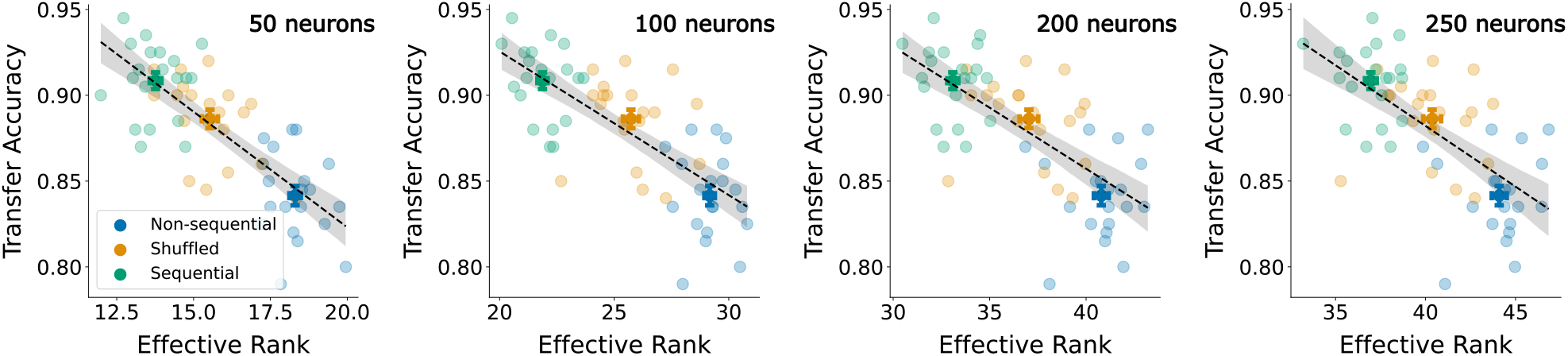
Effect of dimensionality on transfer accuracy is invariant to readout subspace size. Transfer accuracy versus effective rank for different subspace sizes (from left to right: 50 (*r*=-0.78, *p*<<0.0001), 100 (*r*=-0.74, *p*<<0.0001), 200 (*r*=-0.70, *p*<<0.0001), and 250 (*r*=-0.67, *p*<<0.0001) neurons).

**S6 Fig.**
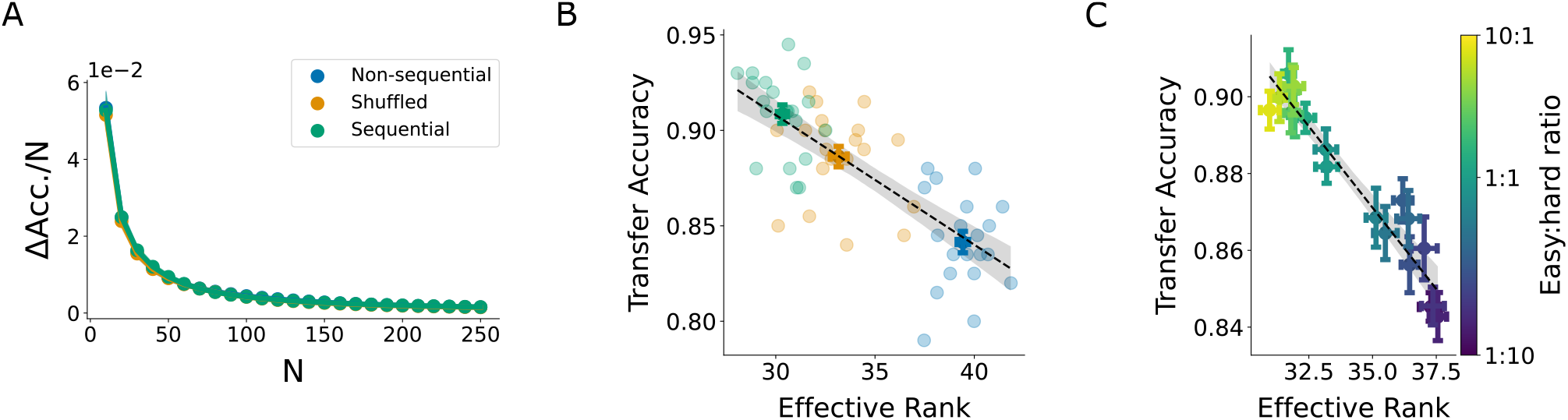
Readout subspace dimensionality resulting from choosing 150 neurons with the largest contribution to the transfer accuracy. **(A)** Accuracy drop per neuron lesioned, plotted against the number of neurons lesioned. The neurons were chosen by their functional contribution to the transfer accuracy. The accuracy drop per neuron plateaued at about N=150, meaning that the readout subspace was mostly affected by about 150 neurons. **(B)** Transfer accuracy versus effective rank for models across curricula (*r*=-0.76, *p*<<0.0001). **(C)** Negative correlation of the effective rank for shuffled curricula with different easy:hard sample ratios with transfer accuracy (*r*=-0.96, *p*<<0.0001).

**S7 Fig.**
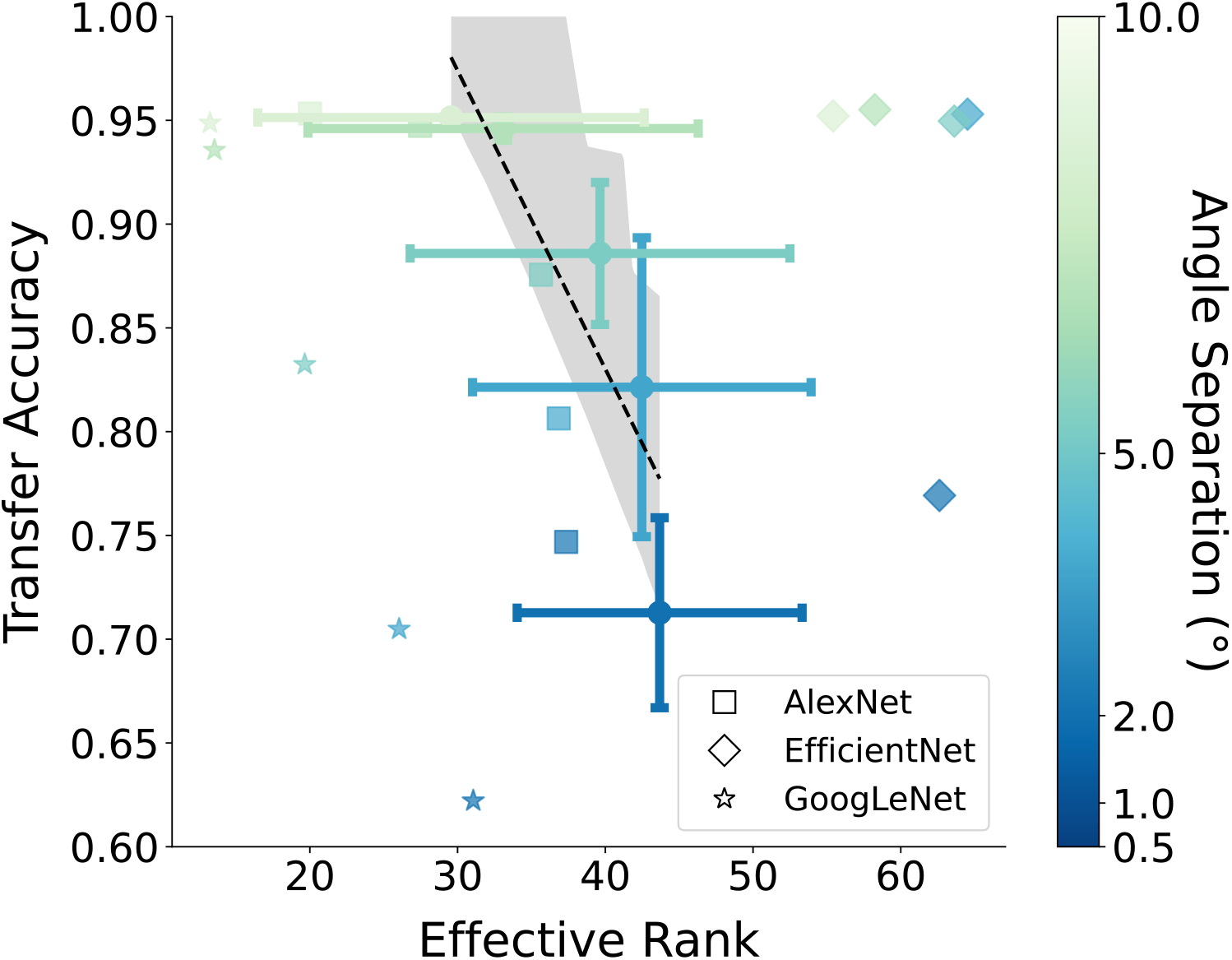
Comparing readout subspace dimensionality in different model backbones. Transfer accuracy versus effective rank for selected model backbones (AlexNet, EfficientNet, and GoogLeNet) trained and tested on single angle separations. Correlation is calculated based on the means across all models for single angle separation (*r*=-0.89, *p*=0.046).

## S1 Appendix. Results for different model backbones

To investigate the robustness of our results across different model backbone choices, we replaced our AlexNet backbone with two other ANN model architectures: GoogLeNet [1] and EfficientNet [2] with added skip connections from each layer to the readout. These models were selected because they could solve the task without extensive hyperparameter engineering and optimization. Additionally, like AlexNet, these models are small convolutional backbones without complex information pathways or spatial averaging. We found that these models show similar patterns of transfer accuracy versus task difficulty, and that the transfer accuracy is negatively correlated with the dimensionality of the readout subspace (Fig. S7; *r*=-0.89, *p*=0.046). However, our results did not extend to other larger and more complex model backbones (see Methods for a complete list of architectures), which are less accurate approximations of single-neuron tuning properties in the visual system [3]. As we showed in Fig. 5, single-neuron tuning properties are key to the observed relationship between dimensionality and generalization, so we conjecture that the models with more similar single-neuron tunings to the brain should better reflect learning generalization in humans. Another potential reason for this discrepancy relates to the preserved fine orientation information required at the output of the model due to the nature of the task. In most of the pretrained convolutional architectures, particularly the deeper models, the fine orientation information diminishes at the output due to spatial averaging through the model. Adding skip connections from every layer to the output could partially solve this problem, but for larger models this makes training intractable without extensive engineering and hyperparameter optimization.

## References

1. Ahissar M. Perceptual Learning. Current Directions in Psychological Science. 1999;8: 124–128. 10.1111/1467-8721.00029

2. Ahissar M, Hochstein S. Learning Pop-out Detection: Specificities to Stimulus Characteristics. Vision Research. 1996;36: 3487–3500. 10.1016/0042-6989(96)00036-3

3. Ahissar M, Hochstein S. Task difficulty and the specificity of perceptual learning. Nature. 1997;387: 401–406. 10.1038/387401a0

4. Fahle M. Perceptual learning: specificity versus generalization. Current Opinion in Neurobiology. 2005;15: 154–160. 10.1016/j.conb.2005.03.010

5. Ahissar M, Hochstein S. The spread of attention and learning in feature search: effects of target distribution and task difficulty. Vision Research. 2000;40: 1349–1364. 10.1016/S0042-6989(00)00002-X

6. Bakhtiari S, Awada A, Pack CC. Influence of stimulus complexity on the specificity of visual perceptual learning. Journal of Vision. 2020;20: 13–13. 10.1167/jov.20.6.13

7. Wang R, Zhang J-Y, Klein SA, Levi DM, Yu C. Vernier perceptual learning transfers to completely untrained retinal locations after double training: A “piggybacking” effect. Journal of Vision. 2014;14: 12–12. 10.1167/14.13.12

8. Xiao L-Q, Zhang J-Y, Wang R, Klein SA, Levi DM, Yu C. Complete transfer of perceptual learning across retinal locations enabled by double training. Curr Biol. 2008;18: 1922–1926. 10.1016/j.cub.2008.10.030

9. Szpiro SFA, Wright BA, Carrasco M. Learning one task by interleaving practice with another task. Vision Research. 2014;101: 118–124. 10.1016/j.visres.2014.06.004

10. Hochstein S, Ahissar M. View from the Top: Hierarchies and Reverse Hierarchies in the Visual System. Neuron. 2002;36: 791–804. 10.1016/S0896-6273(02)01091-7

11. Yamins DLK, Hong H, Cadieu CF, Solomon EA, Seibert D, DiCarlo JJ. Performance-optimized hierarchical models predict neural responses in higher visual cortex. Proceedings of the National Academy of Sciences. 2014;111: 8619–8624. 10.1073/pnas.1403112111

12. Khaligh-Razavi S-M, Kriegeskorte N. Deep Supervised, but Not Unsupervised, Models May Explain IT Cortical Representation. PLOS Computational Biology. 2014;10: 1–29. 10.1371/journal.pcbi.1003915

13. Mineault P, Bakhtiari S, Richards B, Pack C. Your head is there to move you around: Goal-driven models of the primate dorsal pathway. In: Ranzato M, Beygelzimer A, Dauphin Y, Liang PS, Vaughan JW, editors. Advances in Neural Information Processing Systems. Curran Associates, Inc.; 2021. pp. 28757–28771. Available: https://proceedings.neurips.cc/paper_files/paper/2021/file/f1676935f9304b97d59b0738289d2e22-Paper.pdf

14. Bakhtiari S, Mineault P, Lillicrap T, Pack C, Richards B. The functional specialization of visual cortex emerges from training parallel pathways with self-supervised predictive learning. In: Ranzato M, Beygelzimer A, Dauphin Y, Liang PS, Vaughan JW, editors. Advances in Neural Information Processing Systems. Curran Associates, Inc.; 2021. pp. 25164–25178. Available: https://proceedings.neurips.cc/paper_files/paper/2021/file/d384dec9f5f7a64a36b5c8f03b8a6d92-Paper.pdf

15. Schrimpf M, Kubilius J, Hong H, Majaj NJ, Rajalingham R, Issa EB, et al. Brain-Score: Which Artificial Neural Network for Object Recognition is most Brain-Like? bioRxiv. 2020. 10.1101/407007

16. Geirhos R, Narayanappa K, Mitzkus B, Thieringer T, Bethge M, Wichmann FA, et al. Partial success in closing the gap between human and machine vision. In: Ranzato M, Beygelzimer A, Dauphin Y, Liang PS, Vaughan JW, editors. Advances in Neural Information Processing Systems. Curran Associates, Inc.; 2021. pp. 23885–23899. Available: https://proceedings.neurips.cc/paper_files/paper/2021/file/c8877cff22082a16395a57e97232bb6f-Paper.pdf

17. Wenliang LK, Seitz AR. Deep Neural Networks for Modeling Visual Perceptual Learning. Journal of Neuroscience. 2018;38: 6028–6044. 10.1523/JNEUROSCI.1620-17.2018

18. Jeter PE, Dosher BA, Petrov A, Lu ZL. Task precision at transfer determines specificity of perceptual learning. Journal of Vision. 2009;9: 1–1. 10.1167/9.3.1

19. Valois RLD, Albrecht DG, Thorell LG. Spatial frequency selectivity of cells in macaque visual cortex. Vision Research. 1982;22: 545–559. 10.1016/0042-6989(82)90113-4

20. Broderick WF, Simoncelli EP, Winawer J. Mapping spatial frequency preferences across human primary visual cortex. Journal of Vision. 2022;22: 3–3. 10.1167/jov.22.4.3

21. Krizhevsky A, Sutskever I, Hinton GE. ImageNet Classification with Deep Convolutional Neural Networks. In: Pereira F, Burges CJ, Bottou L, Weinberger KQ, editors. Advances in Neural Information Processing Systems. Curran Associates, Inc.; 2012. Available: https://proceedings.neurips.cc/paper_files/paper/2012/file/c399862d3b9d6b76c8436e924a68c45b-Paper.pdf

22. J. Deng, W. Dong, R. Socher, L.-J. Li, Kai Li, Li Fei-Fei. ImageNet: A large-scale hierarchical image database. 2009 IEEE Conference on Computer Vision and Pattern Recognition. 2009. pp. 248–255. 10.1109/CVPR.2009.5206848

23. Cadieu CF, Hong H, Yamins DLK, Pinto N, Ardila D, Solomon EA, et al. Deep Neural Networks Rival the Representation of Primate IT Cortex for Core Visual Object Recognition. PLOS Computational Biology. 2014;10: 1–18. 10.1371/journal.pcbi.1003963

24. Khosla M, Williams AH, McDermott J, Kanwisher N. Privileged representational axes in biological and artificial neural networks. bioRxiv. 2024. 10.1101/2024.06.20.599957

25. Bakhtiari S. Can Deep Learning Model Perceptual Learning? Journal of Neuroscience. 2019;39: 194–196. 10.1523/JNEUROSCI.2209-18.2018

26. Liu LD, Pack CC. The Contribution of Area MT to Visual Motion Perception Depends on Training. Neuron. 2017;95: 436–446.e3. 10.1016/j.neuron.2017.06.024

27. Chowdhury SA, DeAngelis GC. Fine Discrimination Training Alters the Causal Contribution of Macaque Area MT to Depth Perception. Neuron. 2008;60: 367–377. 10.1016/j.neuron.2008.08.023

28. Law C-T, Gold JI. Neural correlates of perceptual learning in a sensory-motor, but not a sensory, cortical area. Nature Neuroscience. 2008;11: 505–513. 10.1038/nn2070

29. Grunewald A, Linden JF, Andersen RA. Responses to Auditory Stimuli in Macaque Lateral Intraparietal Area I. Effects of Training. Journal of Neurophysiology. 1999;82: 330–342. 10.1152/jn.1999.82.1.330

30. Dosher BA, Jeter P, Liu J, Lu Z-L. An integrated reweighting theory of perceptual learning. Proceedings of the National Academy of Sciences. 2013;110: 13678–13683. 10.1073/pnas.1312552110

31. Wu X, Dyer E, Neyshabur B. When Do Curricula Work? International Conference on Learning Representations. 2021. Available: https://openreview.net/forum?id=tW4QEInpni

32. Liu Z, Liu Y, Michaud EJ, Gore J, Tegmark M. Physics of Skill Learning. 2025. Available: https://arxiv.org/abs/2501.12391

33. Bengio Y, Louradour J, Collobert R, Weston J. Curriculum learning. Proceedings of the 26th Annual International Conference on Machine Learning. New York, NY, USA: Association for Computing Machinery; 2009. pp. 41–48. 10.1145/1553374.1553380

34. Hacohen G, Weinshall D. On The Power of Curriculum Learning in Training Deep Networks. In: Chaudhuri K, Salakhutdinov R, editors. Proceedings of the 36th International Conference on Machine Learning. PMLR; 2019. pp. 2535–2544. Available: https://proceedings.mlr.press/v97/hacohen19a.html

35. Roy O, Vetterli M. The effective rank: A measure of effective dimensionality. 2007 15th European Signal Processing Conference. 2007. pp. 606–610. Available: https://www.eurasip.org/Proceedings/Eusipco/Eusipco2007/Papers/a5p-h05.pdf

36. Elmoznino E, Bonner MF. High-performing neural network models of visual cortex benefit from high latent dimensionality. PLOS Computational Biology. 2024;20: 1–23. 10.1371/journal.pcbi.1011792

37. Mazurek M, Kager M, Van Hooser SD. Robust quantification of orientation selectivity and direction selectivity. Frontiers in Neural Circuits. 2014;Volume 8–2014. 10.3389/fncir.2014.00092

38. Fine I, Jacobs RA. Comparing perceptual learning across tasks: A review. Journal of Vision. 2002;2: 5–5. 10.1167/2.2.5

39. Hung S-C, Seitz AR. Prolonged Training at Threshold Promotes Robust Retinotopic Specificity in Perceptual Learning. J Neurosci. 2014;34: 8423. 10.1523/JNEUROSCI.0745-14.2014

40. Petrov AA, Dosher BA, Lu Z-L. The dynamics of perceptual learning: an incremental reweighting model. Psychol Rev. 2005;112: 715–743. 10.1037/0033-295X.112.4.715

41. Karni A, Sagi D. Where practice makes perfect in texture discrimination: evidence for primary visual cortex plasticity. Proceedings of the National Academy of Sciences. 1991;88: 4966–4970. 10.1073/pnas.88.11.4966

42. Kawaguchi K, Bengio Y, Kaelbling L. Generalization in Deep Learning. In: Grohs P, Kutyniok G, editors. Mathematical Aspects of Deep Learning. Cambridge: Cambridge University Press; 2022. pp. 112–148. 10.1017/9781009025096.003

43. Canatar A, Bordelon B, Pehlevan C. Spectral bias and task-model alignment explain generalization in kernel regression and infinitely wide neural networks. Nature Communications. 2021;12: 2914. 10.1038/s41467-021-23103-1

44. Lampinen AK, Ganguli S. An analytic theory of generalization dynamics and transfer learning in deep linear networks. International Conference on Learning Representations. 2019. Available: https://openreview.net/forum?id=ryfMLoCqtQ

45. Fel T, Béthune L, Lampinen AK, Serre T, Hermann K. Understanding Visual Feature Reliance through the Lens of Complexity. In: Globerson A, Mackey L, Belgrave D, Fan A, Paquet U, Tomczak J, et al., editors. Advances in Neural Information Processing Systems. Curran Associates, Inc.; 2024. pp. 69888–69924. Available: https://proceedings.neurips.cc/paper_files/paper/2024/file/819977c0a95458911bbfd9e5b5115018-Paper-Conference.pdf

46. Semedo JD, Zandvakili A, Machens CK, Yu BM, Kohn A. Cortical Areas Interact through a Communication Subspace. Neuron. 2019;102: 249–259.e4. 10.1016/j.neuron.2019.01.026

47. Kohn A, Jasper AI, Semedo JD, Gokcen E, Machens CK, Yu BM. Principles of Corticocortical Communication: Proposed Schemes and Design Considerations. Trends in Neurosciences. 2020;43: 725–737. 10.1016/j.tins.2020.07.001

48. MacDowell CJ, Libby A, Jahn CI, Tafazoli S, Ardalan A, Buschman TJ. Multiplexed subspaces route neural activity across brain-wide networks. Nature Communications. 2025;16: 3359. 10.1038/s41467-025-58698-2

49. Bounds HA, Adesnik H. Network influence determines the impact of cortical ensembles on stimulus detection. Neuron. 2025;113: 2358–2369.e5. 10.1016/j.neuron.2025.04.023

50. Elman JL. Learning and development in neural networks: the importance of starting small. Cognition. 1993;48: 71–99. 10.1016/0010-0277(93)90058-4

51. Pentina A, Sharmanska V, Lampert CH. Curriculum learning of multiple tasks. 2015 IEEE Conference on Computer Vision and Pattern Recognition (CVPR). 2015. pp. 5492–5500. 10.1109/CVPR.2015.7299188

52. Stretcu O, Platanios EA, Mitchell TM, Póczos B. Coarse-to-Fine Curriculum Learning. 2021. Available: https://arxiv.org/abs/2106.04072

53. Beukers AO, Collin SHP, Kempner RP, Franklin NT, Gershman SJ, Norman KA. Blocked training facilitates learning of multiple schemas. Communications Psychology. 2024;2: 28. 10.1038/s44271-024-00079-4

54. Surkov M, Mosin V, Yamshchikov I. Do Data-based Curricula Work? Proceedings of the Third Workshop on Insights from Negative Results in NLP. Association for Computational Linguistics; 2022. pp. 119–128. 10.18653/v1/2022.insights-1.16

55. Carlini N, Erlingsson Ú, Papernot N. Distribution Density, Tails, and Outliers in Machine Learning: Metrics and Applications. 2019. Available: https://arxiv.org/abs/1910.13427

56. Saglietti L, Mannelli S, Saxe A. An Analytical Theory of Curriculum Learning in Teacher-Student Networks. In: Koyejo S, Mohamed S, Agarwal A, Belgrave D, Cho K, Oh A, editors. Advances in Neural Information Processing Systems. Curran Associates, Inc.; 2022. pp. 21113–21127. Available: https://proceedings.neurips.cc/paper_files/paper/2022/file/84bad835faaf48f24d990072bb5b80ee-Paper-Conference.pdf

57. Gong T, Zhao Q, Meng D, Xu Z. Why curriculum learning & self-paced learning work in big/noisy data: A theoretical perspective. Big Data & Information Analytics. 2015;1: 111–127. 10.3934/bdia.2016.1.111

58. Zhang Y, Mohamed A, Abdine H, Shang G, Vazirgiannis M. Beyond Random Sampling: Efficient Language Model Pretraining via Curriculum Learning. 2025. Available: https://arxiv.org/abs/2506.11300

59. Nagatsuka K, Broni-Bediako C, Atsumi M. Length-Based Curriculum Learning for Efficient Pre-training of Language Models. New Generation Computing. 2023;41: 109–134. 10.1007/s00354-022-00198-8

60. Chang E, Yeh H-S, Demberg V. Does the Order of Training Samples Matter? Improving Neural Data-to-Text Generation with Curriculum Learning. In: Merlo P, Tiedemann J, Tsarfaty R, editors. Online: Association for Computational Linguistics; 2021. pp. 727–733. 10.18653/v1/2021.eacl-main.61

61. Hocker D, Constantinople CM, Savin C. Compositional pretraining improves computational efficiency and matches animal behaviour on complex tasks. Nature Machine Intelligence. 2025;7: 689–702. 10.1038/s42256-025-01029-3

62. Cohn D, Ghahramani Z, Jordan M. Active Learning with Statistical Models. In: Tesauro G, Touretzky D, Leen T, editors. Advances in Neural Information Processing Systems. MIT Press; 1994. Available: https://proceedings.neurips.cc/paper_files/paper/1994/file/7f975a56c761db6506eca0b37ce6ec87-Paper.pdf

63. Yang J, Yan F-F, Chen L, Xi J, Fan S, Zhang P, et al. General learning ability in perceptual learning. Proceedings of the National Academy of Sciences. 2020;117: 19092–19100. 10.1073/pnas.2002903117

64. Holton E, Braun L, Thompson JAF, Grohn J, Summerfield C. Humans and neural networks show similar patterns of transfer and interference during continual learning. PsyArXiv; 2025. 10.31234/osf.io/98ksw_v1

65. Liebana S, Laffere A, Toschi C, Schilling L, Moretti J, Podlaski J, et al. Dopamine encodes deep network teaching signals for individual learning trajectories. Cell. 2025;188: 3789–3805. 10.1016/j.cell.2025.05.025

66. Pagan M, Tang VD, Aoi MC, Pillow JW, Mante V, Sussillo D, et al. Individual variability of neural computations underlying flexible decisions. Nature. 2025;639: 421–429. 10.1038/s41586-024-08433-6

67. Laamerad P, Awada A, Pack CC, Bakhtiari S. Asymmetric stimulus representations bias visual perceptual learning. Journal of Vision. 2024;24: 10–10. 10.1167/jov.24.1.10

68. Bordelon B, Pehlevan C. Population codes enable learning from few examples by shaping inductive bias. eLife. 2022;11: e78606. 10.7554/eLife.78606

69. Bordelon B, Canatar A, Pehlevan C. Spectrum Dependent Learning Curves in Kernel Regression and Wide Neural Networks. In: Iii HD, Singh A, editors. Proceedings of Machine Learning Research. PMLR; 2020. pp. 1024–1034. Available: https://proceedings.mlr.press/v119/bordelon20a.html

70. Jin M, Beck JM, Glickfeld LL. Neuronal Adaptation Reveals a Suboptimal Decoding of Orientation Tuned Populations in the Mouse Visual Cortex. J Neurosci. 2019;39: 3867. 10.1523/JNEUROSCI.3172-18.2019

71. Laamerad P, Krause MR, Guitton D, Pack CC. Inactivation of primate cortex reveals inductive biases in visual learning. Current Biology. 2025;35: 4699–4713. 10.1016/j.cub.2025.08.027

72. Driscoll LN, Shenoy K, Sussillo D. Flexible multitask computation in recurrent networks utilizes shared dynamical motifs. Nature Neuroscience. 2024;27: 1349–1363. 10.1038/s41593-024-01668-6

73. Panigrahi A, Liu B, Malladi S, Risteski A, Goel S. Progressive distillation induces an implicit curriculum. International Conference on Learning Representations. 2025. Available: https://openreview.net/forum?id=wPMRwmytZe

74. Seitz AR. Perceptual learning. Current Biology. 2017;27: R631–R636.

75. Waite S, Grigorian A, Alexander RG, Macknik SL, Carrasco M, Heeger DJ, et al. Analysis of Perceptual Expertise in Radiology - Current Knowledge and a New Perspective. Front Hum Neurosci. 2019;13: 213. https://doi.org/0.3389/fnhum.2019.00213

76. Hornsby AN, Love BC. Improved Classification of Mammograms Following Idealized Training. J Appl Res Mem Cogn. 2014;3: 72–76. 10.1016/j.jarmac.2014.04.009

77. Roads BD, Xu B, Robinson JK, Tanaka JW. The easy-to-hard training advantage with real-world medical images. Cogn Res Princ Implic. 2018;3: 38. 10.1186/s41235-018-0131-6

78. Rosner J. The Development and Validation of an Individualized Perceptual Skills Curriculum. 1972. Available: https://api.semanticscholar.org/CorpusID:141790835

79. Rau MA, Aleven V, Rummel N. Interleaved practice in multi-dimensional learning tasks: Which dimension should we interleave? Learning and Instruction. 2013;23: 98–114. 10.1016/j.learninstruc.2012.07.003

80. Chen W, Chen C, Li B, Zhang J. Applying Interleaving Strategy of Learning Materials and Perceptual Modality to Address Secondary Students’ Need to Restore Cognitive Capacity. International Journal of Environmental Research and Public Health. 2022;19. 10.3390/ijerph19127505

81. Samani J, Pan SC. Interleaved practice enhances memory and problem-solving ability in undergraduate physics. npj Science of Learning. 2021;6: 32. 10.1038/s41539-021-00110-x

82. Jiang L, Meng D, Zhao Q, Shan S, Hauptmann A. Self-Paced Curriculum Learning. Proceedings of the AAAI Conference on Artificial Intelligence. 2015;29. 10.1609/aaai.v29i1.9608

83. Noh SM, Yan VX, Bjork RA, Maddox WT. Optimal sequencing during category learning: Testing a dual-learning systems perspective. Cognition. 2016;155: 23–29. 10.1016/j.cognition.2016.06.007

84. Russin J, Pavlick E, Frank MJ. Parallel trade-offs in human cognition and neural networks: The dynamic interplay between in-context and in-weight learning. Proceedings of the National Academy of Sciences. 2025;122: e2510270122. 10.1073/pnas.2510270122

85. Pesnot Lerousseau J, Summerfield C. Shared sensitivity to data distribution during learning in humans and transformer networks. Nature Human Behaviour. 2026;10: 601–614. 10.1038/s41562-025-02359-3

86. Whittington JCR, Bogacz R. Theories of Error Back-Propagation in the Brain. Trends Cogn Sci. 2019;23: 235–250. 10.1016/j.tics.2018.12.005

87. Lillicrap TP, Santoro A, Marris L, Akerman CJ, Hinton G. Backpropagation and the brain. Nature Reviews Neuroscience. 2020;21: 335–346. 10.1038/s41583-020-0277-3

88. Cornford J, Pogodin R, Ghosh A, Sheng K, Bicknell BA, Codol O, et al. Brain-like learning with exponentiated gradients. bioRxiv. 2024. 10.1101/2024.10.25.620272

89. Ororbia A, Mali A, Kohan A, Millidge B, Salvatori T. A Review of Neuroscience-Inspired Machine Learning. arXiv [csNE]. 2024. Available: http://arxiv.org/abs/2403.18929

90. Shen S, Sun Y, Lu J, Li C, Chen Q, Mo C, et al. Profiles of visual perceptual learning in feature space. iScience. 2024;27: 109128. 10.1016/j.isci.2024.109128

91. Manenti GL, Dizaji AS, Schwiedrzik CM. Variability in training unlocks generalization in visual perceptual learning through invariant representations. Current Biology. 2023;33: 817–826.e3. 10.1016/j.cub.2023.01.011

92. Peirce JW. PsychoPy—Psychophysics software in Python. Journal of Neuroscience Methods. 2007;162: 8–13. 10.1016/j.jneumeth.2006.11.017

93. Russakovsky O, Deng J, Su H, Krause J, Satheesh S, Ma S, et al. ImageNet Large Scale Visual Recognition Challenge. CoRR. 2014;abs/1409.0575. Available: http://arxiv.org/abs/1409.0575

## Supplementary references

1. Szegedy C, Liu W, Jia Y, Sermanet P, Reed S, Anguelov D, et al. Going Deeper with Convolutions. Computer Vision and Pattern Recognition (CVPR). 2015. Available: http://arxiv.org/abs/1409.4842

2. Tan M, Le Q. EfficientNet: Rethinking Model Scaling for Convolutional Neural Networks. In: Chaudhuri K, Salakhutdinov R, editors. Proceedings of the 36th International Conference on Machine Learning. PMLR; 2019. pp. 6105–6114. Available: https://proceedings.mlr.press/v97/tan19a.html

3. Khosla M, Williams AH, McDermott J, Kanwisher N. Privileged representational axes in biological and artificial neural networks. bioRxiv. 2024. doi:10.1101/2024.06.20.599957

