## Supplementary materials for "The curriculum effect in visual learning: the role of readout dimensionality"

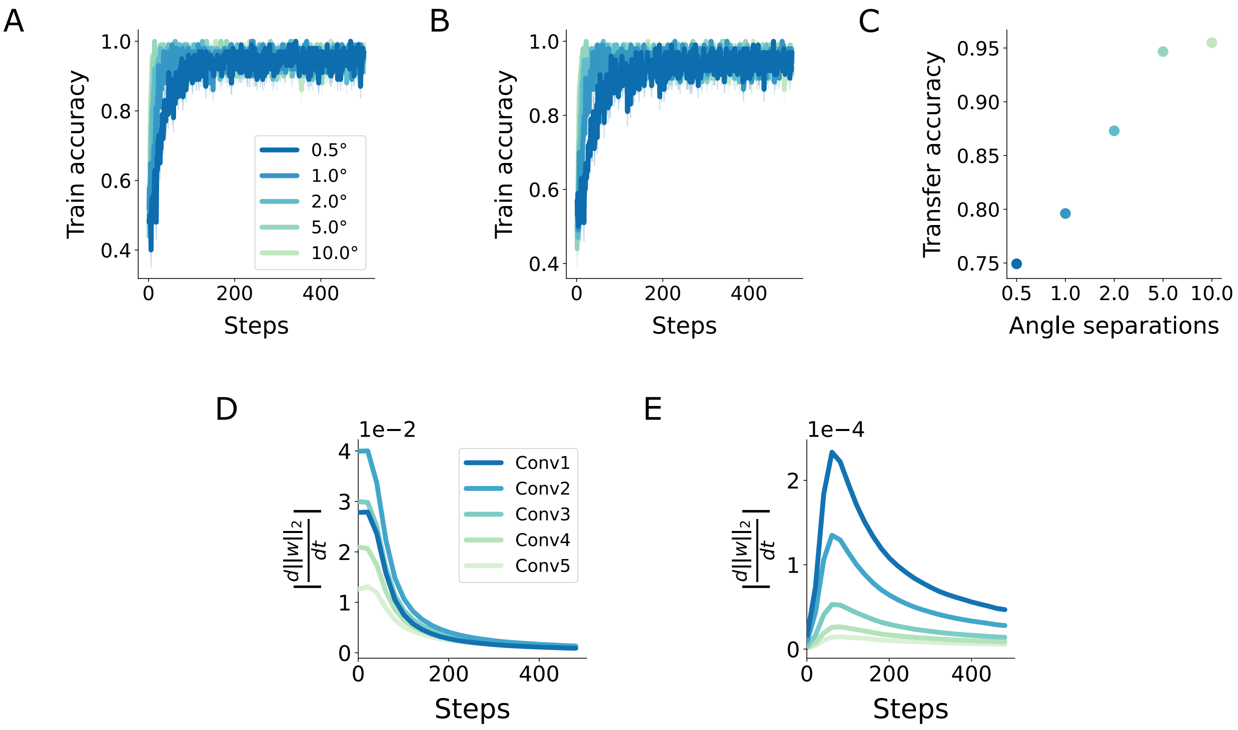


**S1 Fig. Additional training, transfer accuracy, and weight change data.** (**A**) Train accuracy with standard error visible. (**B**) Train accuracy with standard error visible for a model with convolutional and pooling layer weights frozen and readout weights unfrozen. (**C**) Transfer accuracy for the model from B. **(D)** Gradient of the weight norm of skip connection weights throughout training on a 0.5° angle separation. **(E)** Gradient of the weight norm of convolutional weights throughout training on a 0.5° angle separation.

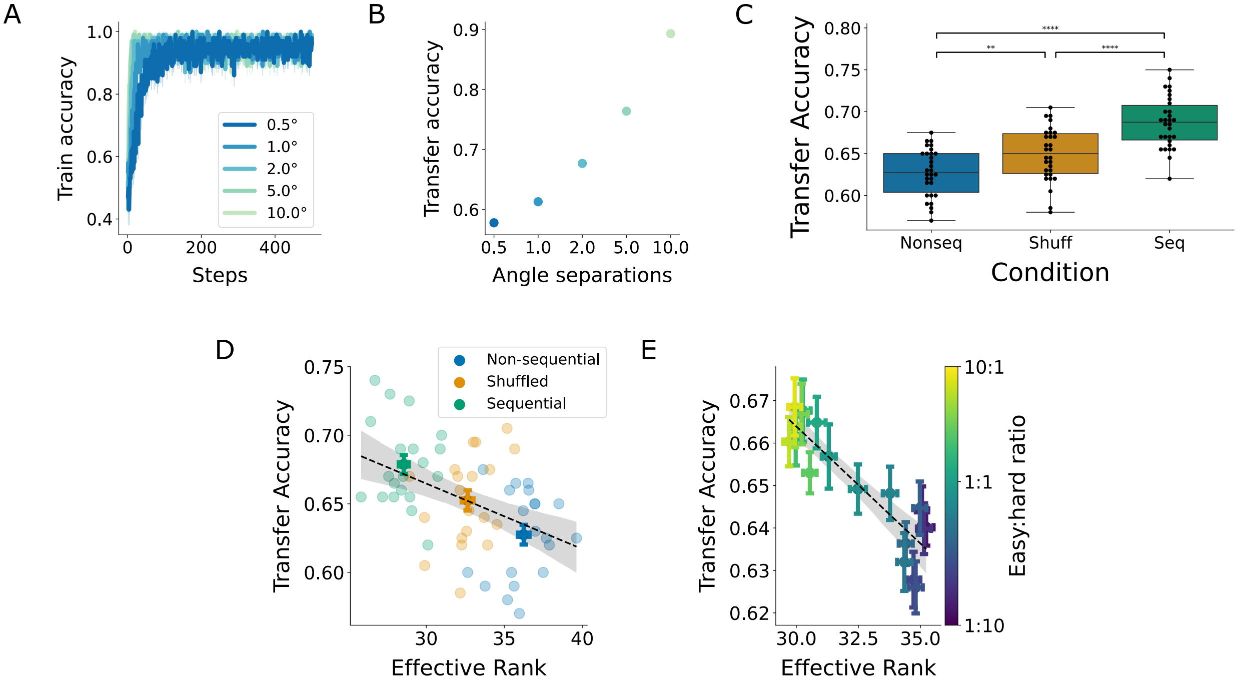


**S2 Fig. Results for a changed reference orientation in the transfer condition.** **(A)** Train accuracy. **(B)** Transfer accuracy. **(C)** Transfer accuracy for models across curricula. **(D)** Transfer accuracy versus effective rank for models across curricula ($r$=-0.45, $p$=0.00034). **(E)** Negative correlation of the effective rank for shuffled curricula with different easy:hard sample ratios with transfer accuracy ($r$=-0.89, $p$<<0.0001).

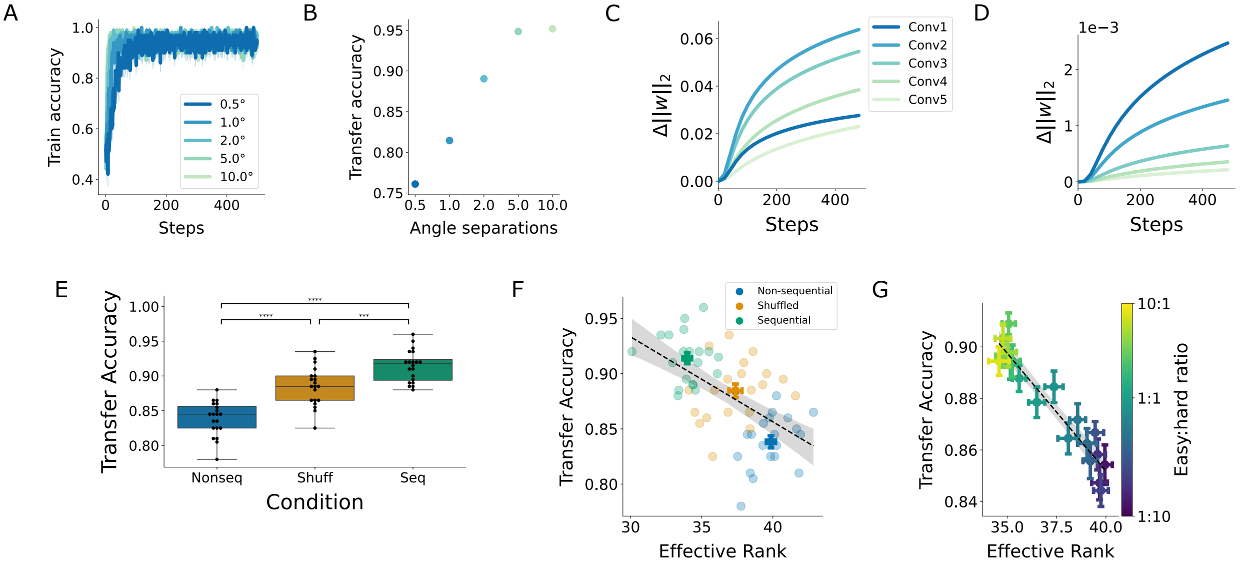


**S3 Fig. Results for models with randomly initialized readout weights.** **(A)** Train accuracy. **(B)** Transfer accuracy. **(C)** Change in the L2-norm of skip connection weights throughout training on a 0.5° angle separation. **(D)** Change in the L2-norm of convolutional weights throughout training on a 0.5° angle separation. **(E)** Transfer accuracy for models across curricula. **(F)** Transfer accuracy versus effective rank for models across curricula ($r$=-0.57, $p$<<0.0001). **(G)** Negative correlation of the effective rank for shuffled curricula with different easy:hard sample ratios with transfer accuracy ($r$=-0.95, $p$<<0.0001).

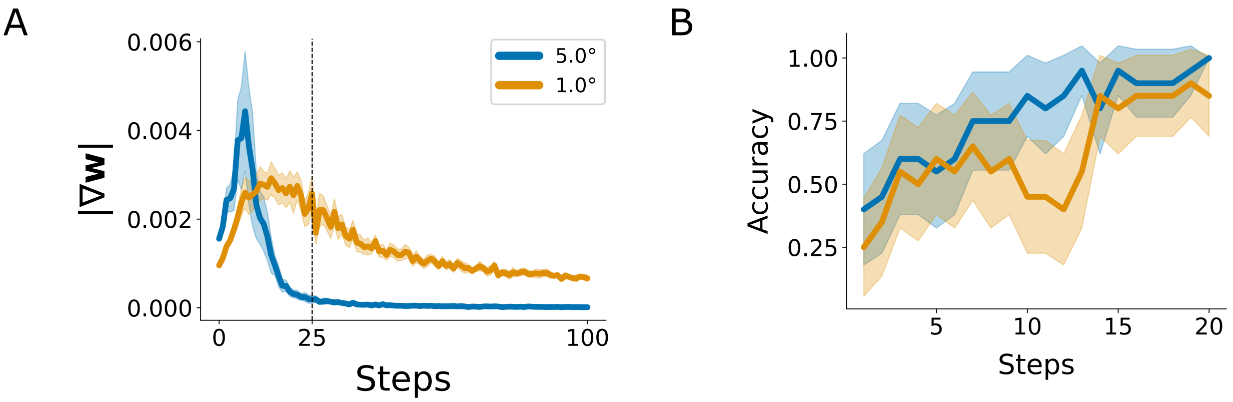


**S4 Fig. Additional data supporting an implicit curriculum in the models. (A)** Parameter gradients for models trained on the shuffled task condition in the first 100 steps of training. This plot includes gradients for the non-active condition at each timestep. **(B)** Accuracy throughout training for models on the shuffled condition shown separately for easy (5°) and hard (1°) samples.

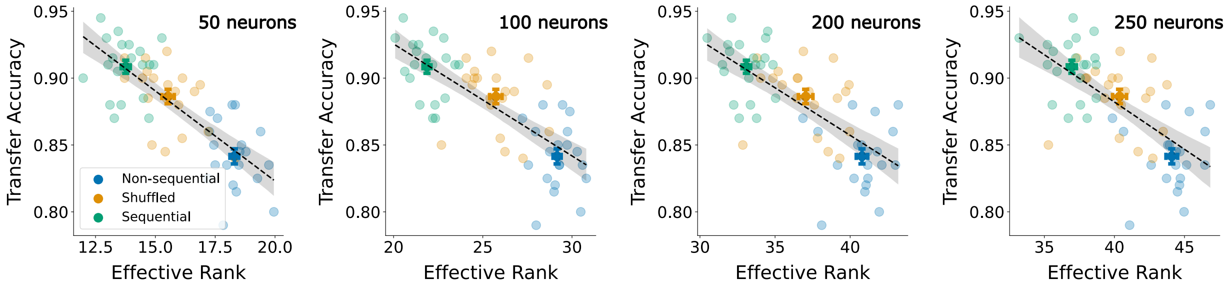


**S5 Fig. Effect of dimensionality on transfer accuracy is invariant to readout subspace size.** Transfer accuracy versus effective rank for different subspace sizes (from left to right: 50 ($r$=-0.78, $p$<<0.0001), 100 ($r$=-0.74, $p$<<0.0001), 200 ($r$=-0.70, $p$<<0.0001), and 250 ($r$=-0.67, $p$<<0.0001) neurons).

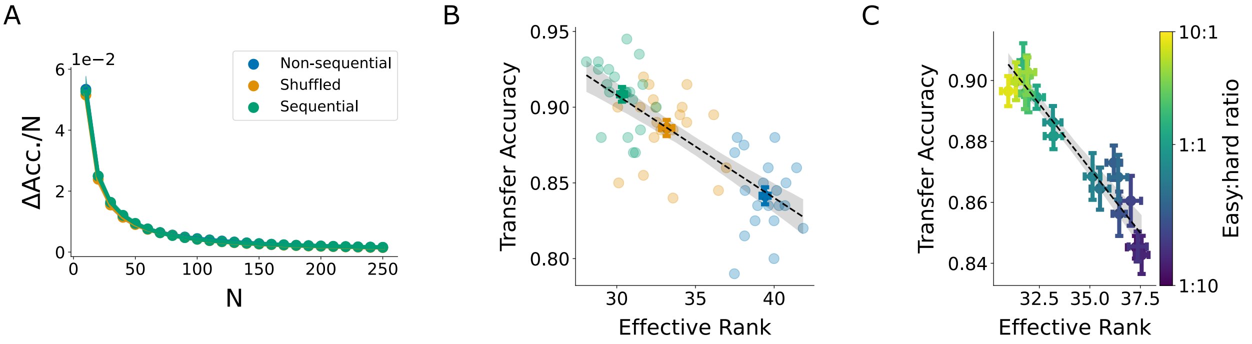


**S6 Fig. Readout subspace dimensionality resulting from choosing 150 neurons with the largest contribution to the transfer accuracy. (A)** Accuracy drop per neuron lesioned, plotted against the number of neurons lesioned. The neurons were chosen by their functional contribution to the transfer accuracy. The accuracy drop per neuron plateaued at about N=150, meaning that the readout subspace was mostly affected by about 150 neurons. **(B)** Transfer accuracy versus effective rank for models across curricula ($r$=-0.76, $p$<<0.0001). **(C)** Negative correlation of the effective rank for shuffled curricula with different easy:hard sample ratios with transfer accuracy ($r$=-0.96, $p$<<0.0001).

**S1 Appendix. Results for different model backbones.**

To investigate the robustness of our results across different model backbone choices, we replaced our AlexNet backbone with two other ANN model architectures: GoogLeNet [1] and EfficientNet [2] with added skip connections from each layer to the readout. These models were selected because they could solve the task without extensive hyperparameter engineering and optimization. Additionally, like AlexNet, these models are small convolutional backbones without complex information pathways or spatial averaging. We found that these models show similar patterns of transfer accuracy versus task difficulty, and that the transfer accuracy is negatively correlated with the dimensionality of the readout subspace (Fig. S7; *r*=-0.89, *p*=0.046). However, our results did not extend to other larger and more complex model backbones (see Methods for a complete list of architectures), which are less accurate approximations of single-neuron tuning properties in the visual system [3]. As we showed in Fig. 5, single-neuron tuning properties are key to the observed relationship between dimensionality and generalization, so we conjecture that the models with more similar single-neuron tunings to the brain should better reflect learning generalization in humans. Another potential reason for this discrepancy relates to the preserved fine orientation information required at the output of the model due to the nature of the task. In most of the pretrained convolutional architectures, particularly the deeper models, the fine orientation information diminishes at the output due to spatial averaging through the model. Adding skip connections from every layer to the output could partially solve this problem, but for larger models this makes training intractable without extensive engineering and hyperparameter optimization.

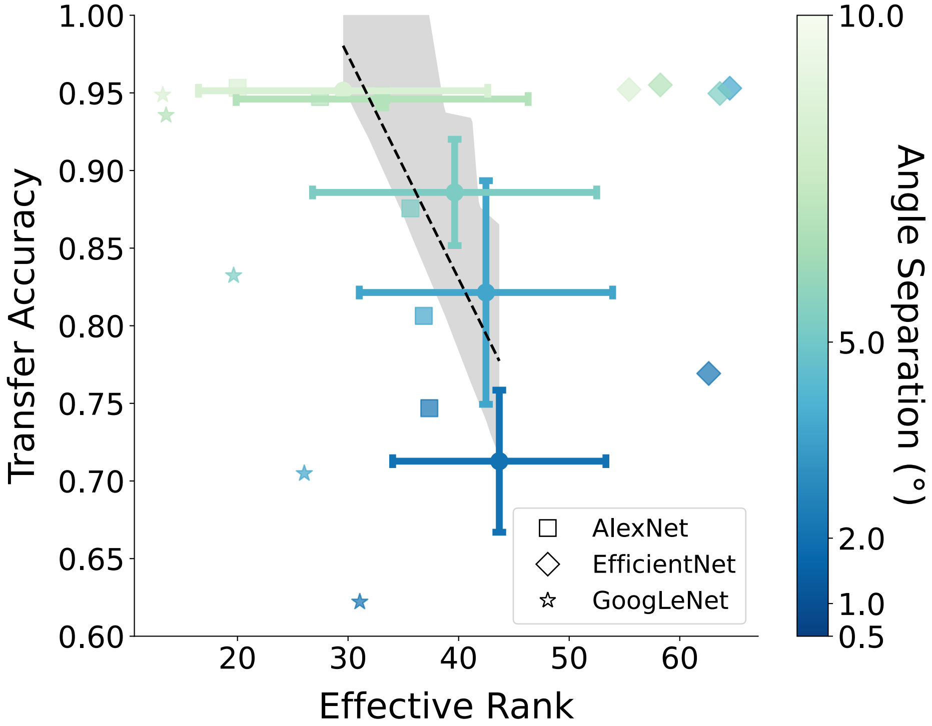


**S7 Fig. Comparing readout subspace dimensionality in different model backbones.** Transfer accuracy versus effective rank for selected model backbones (AlexNet, EfficientNet, and GoogLeNet) trained and tested on single angle separations. Correlation is calculated based on the means across all models for single angle separation (*r*=-0.89, *p*=0.046).

**Supplementary references**

1. Szegedy C, Liu W, Jia Y, Sermanet P, Reed S, Anguelov D, et al. Going Deeper with Convolutions. Computer Vision and Pattern Recognition (CVPR). 2015. Available: http://arxiv.org/abs/1409.4842

2. Tan M, Le Q. EfficientNet: Rethinking Model Scaling for Convolutional Neural Networks. In: Chaudhuri K, Salakhutdinov R, editors. Proceedings of the 36th International Conference on Machine Learning. PMLR; 2019. pp. 6105–6114. Available: https://proceedings.mlr.press/v97/tan19a.html

3. Khosla M, Williams AH, McDermott J, Kanwisher N. Privileged representational axes in biological and artificial neural networks. bioRxiv. 2024. doi:10.1101/2024.06.20.599957
